# Genome-scale mapping of DNA damage suppressors identifies GNB1L as essential for ATM and ATR biogenesis

**DOI:** 10.1101/2022.09.23.508845

**Authors:** Yichao Zhao, Daniel Tabet, Diana Rubio Contreras, Arne Nedergaard Kousholt, Jochen Weile, Henrique Melo, Lisa Hoeg, Atina G. Coté, Zhen-Yuan Lin, Dheva Setiaputra, Jos Jonkers, Anne-Claude Gingras, Fernando Gómez Herreros, Frederick P. Roth, Daniel Durocher

## Abstract

To maintain genome integrity, cells must avoid DNA damage by ensuring the accurate duplication of the genome and by having efficient repair and signaling systems that counteract the genome-destabilizing potential of DNA lesions. To uncover genes and pathways that suppress DNA damage in human cells, we undertook genome-scale CRISPR/Cas9 screens that monitored the levels of DNA damage in the absence or presence of DNA replication stress. We identified 160 genes in RKO cells whose mutation caused high levels of DNA damage in the absence of exogenous genotoxic treatment. This list was highly enriched in essential genes, highlighting the importance of genomic integrity for cellular fitness. Furthermore, the majority of these 160 genes are involved in a limited set of biological processes related to DNA replication and repair, nucleotide biosynthesis, RNA metabolism and iron sulfur cluster biogenesis, suggesting that genome integrity may be insulated from a wide range of cellular processes. Among the many genes identified and validated in this study, we discovered that *GNB1L*, a schizophrenia/autism-susceptibility gene implicated in 22q11.2 syndrome, protects cells from replication catastrophe promoted by mild DNA replication stress. We show that *GNB1L* is involved in the biogenesis of ATR and related phosphatidylinositol 3-kinase-related kinases (PIKKs) through its interaction with the TTT co-chaperone complex. These results implicate PIKK biogenesis as a potential root cause for the neuropsychiatric phenotypes associated with 22q11.2 syndrome. The phenotypic mapping of genes that suppress DNA damage in human cells therefore provides a powerful approach to probe genome maintenance mechanisms.

## Introduction

Perturbed DNA replication is a major source of DNA damage, which has been linked to many human diseases, including cancer (Zeman and Cimprich 2014; Gaillard et al. 2015; Saxena and Zou 2022). During replication, the replisome often encounters impediments that can hinder or block replication fork progression, a phenomenon referred to as DNA replication stress (Zeman and Cimprich 2014). These obstacles include unrepaired DNA lesions, DNA repair intermediates, transcribing RNA polymerases, RNA-DNA hybrids, and non-B DNA structures such as G-quadruplexes (Zeman and Cimprich 2014; Gaillard et al. 2015; Saxena and Zou 2022). Chemical or physical agents can also cause DNA damage that impairs DNA replication. For example, ultraviolet radiation (UV), ionizing radiation (IR) and many genotoxic chemotherapies perturb DNA synthesis.

To minimize the impact of DNA replication perturbations on the stability of the genome, cells have evolved systems that ensure a robust replication process. For example, the stretches of single-stranded DNA (ssDNA) that are produced following the uncoupling of the replicative helicase and DNA polymerases can be sensed by the kinase ataxia telangiectasia and Rad3-related (ATR), which mediates cellular responses to replication stress through the phosphorylation of proteins that include CHK1, RPA, variant histone H2AX and SMARCAL1 (Liu et al. 2000; Ward and Chen 2001; Matsuoka et al. 2007; Couch et al. 2013). ATR is recruited to ssDNA via the ATR-interacting protein ATRIP and is activated either through TOPBP1-dependent or ETAA1-dependent pathways (Kumagai et al. 2006; Bass et al. 2016; Haahr et al. 2016). ATR counteracts replication stress at multiple levels including the stabilization of replication forks, regulation of DNA replication origin firing, by ensuring dNTP availability and by mediating cell cycle checkpoint signaling (Saldivar et al. 2017).

ATR belongs to the family of phosphatidylinositol 3-kinase-related kinases (PIKKs). In mammalian cells, PIKKs have diverse functions in the biology of the DNA damage response (ATR, ATM, DNAPK-cs), cellular metabolism and proliferation (mTOR), nonsense-mediated mRNA decay (SMG1), and transcription control (TRRAP) (Lempiainen and Halazonetis 2009; Baretic and Williams 2014). PIKKs are very sizeable proteins consisting of large HEAT repeat-containing N-termini followed by C-terminal kinase domains. The biogenesis and protein stability of PIKKs depend on the chaperone heat shock protein 90 (HSP90), as well as the TELO2-TTI1-TTI2 (TTT) co-chaperone complex (Takai et al. 2007; Horejsi et al. 2010; Hurov et al. 2010; Takai et al. 2010), and biogenesis of PIKK complexes is known to be essential for genome maintenance (Izumi et al. 2012).

In an effort to identify novel genome maintenance factors, including new participants in the response to DNA replication stress, we undertook the chemogenomic profiling of 27 genotoxic agents using CRISPR/Cas9 genetic screens (Olivieri et al. 2020). The resulting dataset identified new DNA repair factors such as ERCC6L2 and new drug mechanisms of action such as the involvement of TOP2 in the cytotoxicity of the G-quadruplex ligands pyridostatin and CX-5461. However, of the many blind spots associated with such screens, the under-representation of essential genes in the dataset was noticeable. Indeed, the gene *RAD51*, which encodes the essential DNA recombinase, should have been identified as a gene promoting resistance to agents that require a functional homologous recombination (HR) pathway. *RAD51* was not identified in our screens, most likely because the screen readout required single guide (sg) RNA representation in the control (untreated) population. Given that genome stability is an essential cellular process, it is likely that many other cell-essential genes that have roles in genome maintenance were also missed.

To address these shortcomings, we surmised that a screen performed shortly after gene inactivation and that used a readout that relies on a DNA damage-linked phenotype would allow capture of essential genes. We established a phenotypic CRISPR/Cas9 screen pipeline that monitors the level of Ser139-phosphorylated histone H2AX (γ-H2AX) and performed genome-scale screens in two colon epithelial cell lines, either in the absence of treatment or in the presence of the following replication perturbing agents: aphidicolin (Aph), a direct inhibitor of B family DNA polymerases that comprise Pol α, Pol δ, Pol ε and Pol ζ in eukaryotes (Cheng and Kuchta 1993); hydroxyurea (HU), an inhibitor of ribonucleotide reductase, leading to depletion of the cellular pool of dNTPs (Krakoff et al. 1968); and cytarabine (Ara-C), a nucleoside analog that both inhibits ribonucleotide reductase activity and blocks DNA replication elongation (Harrington and Perrino 1995). These screens revealed insights into the processes that protect cells from endogenous DNA damage and identified a number of genes not previously linked to genome maintenance. In particular, we found that the neuropsychiatric disorder candidate gene *GNB1L* protects cells from replication catastrophe under mild replication perturbation (Paylor et al. 2006; Williams et al. 2008; Chen et al. 2012). We found that GNB1L promotes the steady-state levels of biogenesis of ATR, ATM and other related PIKKs in collaboration with the TTT complex.

## Results

### Phenotypic CRISPR/Cas9 screens based on γ-H2AX levels

To probe the processes that prevent the formation of DNA damage independently of their impact on cellular fitness, we established a phenotypic CRISPR/Cas9 screening pipeline based on detecting γ-H2AX by flow cytometry (Huang and Darzynkiewicz 2006) (Fig 1A). The serine 139 residue of histone variant H2AX is quickly phosphorylated in response to DNA damage and replication blockage, making it a widely used DNA damage marker (Rogakou et al. 1998).

**Figure 1.**
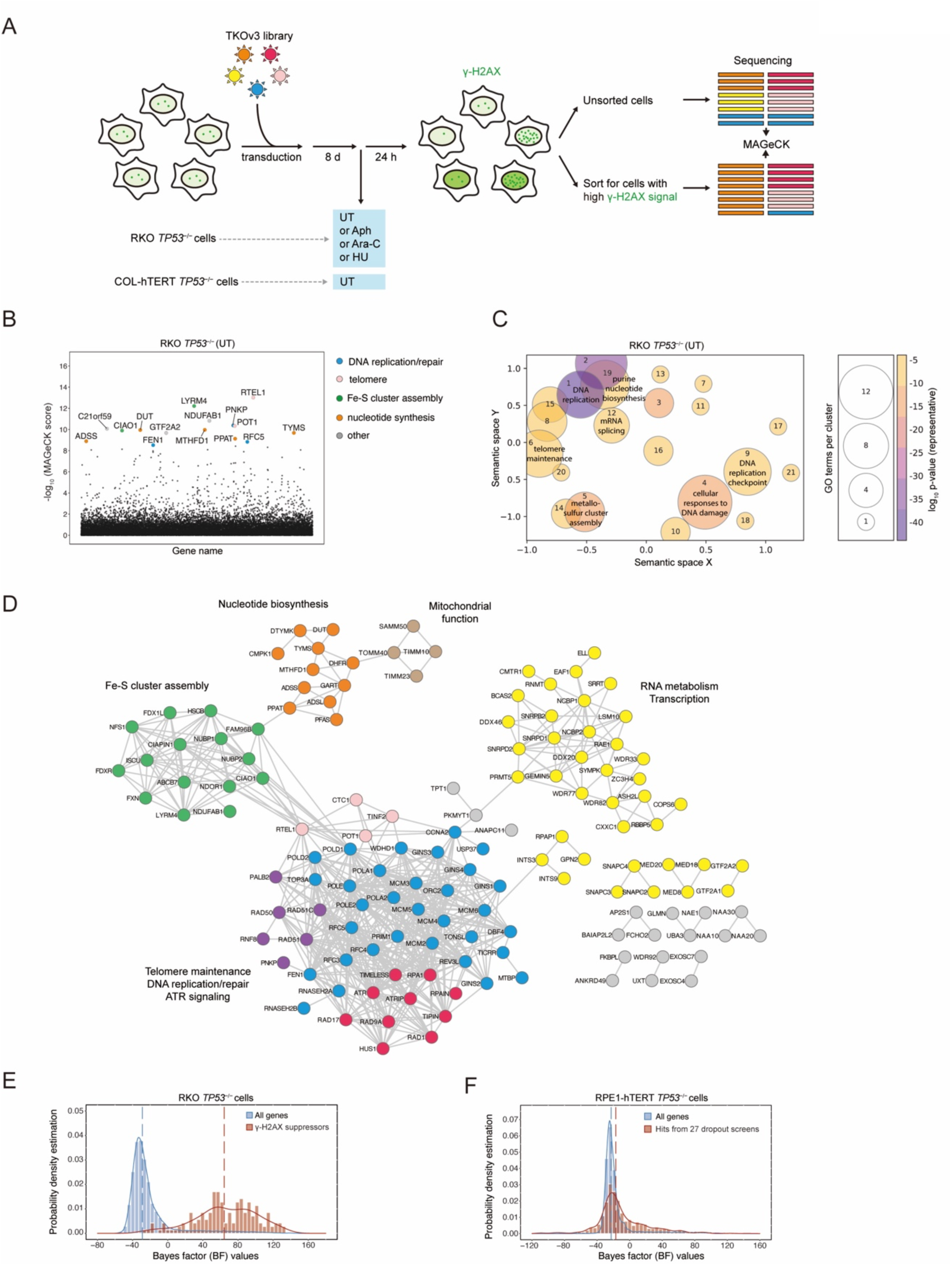
Phenotypic CRISPR screens reveal genes and pathways that prevent endogenous DNA damage in RKO *TP53^−/−^* cells. (A) Schematic of the phenotypic CRISPR screens based on γ-H2AX staining and cell sorting. (B) Manhattan dot plots of the untreated γ-H2AX screen results in untreated (UT) RKO *TP53^−/−^* cells. The top 15 genes are highlighted. (C) Gene Ontology (GO) analysis of Biological Process was performed with Enrichr (https://maayanlab.cloud/Enrichr/) for 160 γ-H2AX suppressors in RKO *TP53^−/−^* cells. The top 80 GO terms ranked by p-value were visualized by GO-Figure! software using a similarity cutoff of 0.5. Color represents the p-value of the representative GO term for the cluster, and the size of the circle represents the number of GO terms in a cluster. (D) STRING network analysis of γ-H2AX suppressors in RKO *TP53^−/−^* cells, based on information from text mining, experiments, databases, and neighborhood, with a high confidence threshold. Out of 160 genes, 138 are mapped to the network. Pathways of genes are manually curated with different colors: green, Fe-S cluster assembly; orange, nucleotide biosynthesis; brown, mitochondrial function; yellow, RNA metabolism and transcription; pink, telomere maintenance; purple, DNA repair; blue, DNA replication; red, ATR signaling; grey, others. (E) Distributions of gene essentiality scores (BF values) of γ-H2AX suppressors (brown) and whole genome reference (blue) in RKO *TP53^−/−^* cells. Kernel density estimation is used for the probability density function. Dashed lines indicate the median for each population. (F) Distributions of gene essentiality scores of hits from 27 dropout genotoxic screens (brown) and whole genome reference (blue) in RPE-hTERT *TP53^−/−^* cells. Dashed lines indicate the median for each population.

We carried out CRISPR/Cas9 screens in the RKO colon carcinoma cell line and in an hTERT-immortalized, i.e. untransformed, colon epithelial cell line, referred to here as COL-hTERT. The *TP53* gene was knocked out by gene editing in both cell lines to prevent the potential confounding effects of p53 activation by genotoxic stress. The cell lines are hereafter referred to as RKO *TP53*^−/−^ and COL-hTERT *TP53*^−/−^ (Fig 1A). We carried out 4 screens in the RKO *TP53*^−/−^ cell line: one screen where cells were left untreated and one screen each in which cells were treated with low doses of Aph, HU and Ara-C. The COL-hTERT *TP53*^−/−^ cell line was only screened in the untreated condition. The screens were carried with the TKOv3 sgRNA library (Hart et al. 2017) and cells with the highest 5% of γ-H2AX fluorescence intensity were sorted. Following sgRNA cassette sequencing, gene-level enrichment scores were computed using MAGeCK comparing sgRNA abundance in the sorted population to that of unsorted cells (Li et al. 2014) (Figs 1A, B, S1A, S2 and Supplementary Table 1).

### Suppressors of γ-H2AX in RKO TP53^−/−^ cells

We identified 160 genes whose mutation caused spontaneous high γ-H2AX levels in RKO *TP53*^−/−^ cells (Supplementary Table 2). This number was obtained by combining 142 genes that scored in the untreated RKO *TP53*^−/−^ screen with a false discovery rate (FDR) value < 0.05 along with any other gene with FDR values between 0.05 to 0.1 that were also a hit (i.e. FDR <0.05) in the drug-treated screens. Gene ontology (GO) analysis of these 160 hits revealed strong enrichment for terms associated with DNA replication (such as GO:0006260), DNA repair (GO:0006281), iron-sulfur cluster metabolism (GO:0016226), DNA damage signaling (GO:0000076), telomere maintenance (GO:0000723), nucleotide metabolism (GO:0009165) and transcription and splicing associated terms (GO:0006366; Fig 1C and Supplementary Table 3). Most of these biological processes are known to promote genome stability, which confirmed the ability of the γ-H2AX phenotypic screens to probe pathways involved in genome maintenance.

Gene-level analysis of the 160 screen hits using STRING (Szklarczyk et al. 2021) mapped 138 genes into a network characterized by 4 connected subnetworks that are enriched in distinct biological processes (Fig 1D). Manual curation of the network nodes revealed a major and dense subcluster of genes with roles in DNA replication, DNA repair, telomere maintenance, and ATR signaling. We also identified subclusters of genes that are involved in nucleotide biosynthesis, RNA metabolism and transcription, as well as iron-sulfur (Fe-S) cluster assembly (Fig 1D).

Inspection of the gene list revealed the presence of many essential genes, such as those encoding multiple components of the CMG helicase responsible for bulk DNA replication (GINS1-4, MCM2-6) and replisome components (POLA1, POLD1, POLE, PRIM1, etc). This observation suggested that the γ-H2AX screens succeeded at identifying essential genes. To explore this possibility more formally, we plotted the Bayes factor (BF) values of the 160 genes obtained in a CRISPR dropout screen in the RKO *TP53*^−/−^ cell line and compared them with the BF score distribution of all genes included in the TKOv3 library. BF values represent the likelihood of gene essentiality, with positive BF values indicating probable essential genes (Hart and Moffat 2016). The distribution of the 160 genes is remarkably shifted to large positive BF values (median value: 64) compared to the median of all genes (−29) (Fig 1E). This is consistent with the notion that genes required to prevent endogenous DNA damage are important for cell survival. In contrast, analysis of the 890 gene hits from our previous fitness-based chemogenomic screens (Olivieri et al. 2020) showed both a distribution and a median BF value similar to that of the library (median values −17 and −22, respectively; Fig 1F). These results indicate that the phenotype-based γ-H2AX screens can better probe the contribution of essential genes to genome maintenance.

### Suppressors of γ-H2AX in COL-hTERT TP53^−/−^ cells

Analysis of the single screen undertaken in the COL-hTERT *TP53^−/−^* cells identified 95 genes whose mutation cause elevated γ-H2AX levels (at FDR < 0.25; Supplementary Table 2). We employed a more relaxed FDR threshold in the COL-hTERT *TP53^−/−^* because of a lower number of replicates done in this screen (see methods). GO analysis and network-based representation using STRING showed a similar set of pathways involved in preventing spontaneous γ-H2AX, including DNA replication (GO:0006260), metallo-sulfur cluster assembly (GO:0031163) and DNA metabolic process (GO:0006260) (Fig S1B, C and Supplementary Table 3). Interestingly, COL-hTERT *TP53^−/−^* cells showed a unique cluster of genes involved in mitochondrial respiration (such as GO: 0042775), suggesting a dependence on this pathway to maintain genome stability. Similarly, analysis of BF values distribution of the hits obtained in the COL-hTERT *TP53^−/−^* screen indicated that the suppressors of spontaneous γ-H2AX formation in this cell line are also enriched in essential genes (Fig S1D).

A total of 32 genes increased γ-H2AX levels in both cell lines when disrupted (Fig S1E, F). Using a hypergeometric distribution, the probability of having 32 genes occurring in both sets by chance alone is 1 x 10^-42^ and therefore this overlap indicates that there is a shared set of genes that prevent DNA damage in both cell lines. Among such genes are those encoding the PRMT5-WDR77 complex, which is actively being pursued as a cancer drug target (Wu et al. 2021), and multiple genes coding proteins involved in nucleotide metabolism and Fe-S cluster assembly (such as DUT, GART, CIAO1 and CIAO2B/FAM96B). Together, these results suggest that there are likely both universal and cell type-restricted genes that prevent spontaneous DNA damage formation, similar to what has been observed with gene essentiality (Hart et al. 2015).

### Validation of genes that prevent endogenous DNA damage formation

To visualize the results of the screens at the gene level, we employed radar plots where gene rank is plotted across the 5 screens. For example, disruption of *CIAO1*, which encodes a WD40 repeat protein involved in Fe-S cluster incorporation (Srinivasan et al. 2007), caused high levels of γ-H2AX in 4 of 5 screen conditions (Fig 2A). In another example, Cas9-mediated disruption of *CFAP298* led to high γ-H2AX levels in 3 of 4 screens performed in RKO *TP53*^−/−^ cells but not in the COL-hTERT *TP53^−/−^* cell screen (Fig 2B). *CFAP298* is an example of a gene not previously linked to genome maintenance. Mutations in *CFAP298* cause primary ciliary dyskinesia, implicating its product in motile cilium function by acting on outer dynein arm assembly (Austin-Tse et al. 2013).

**Figure 2.**
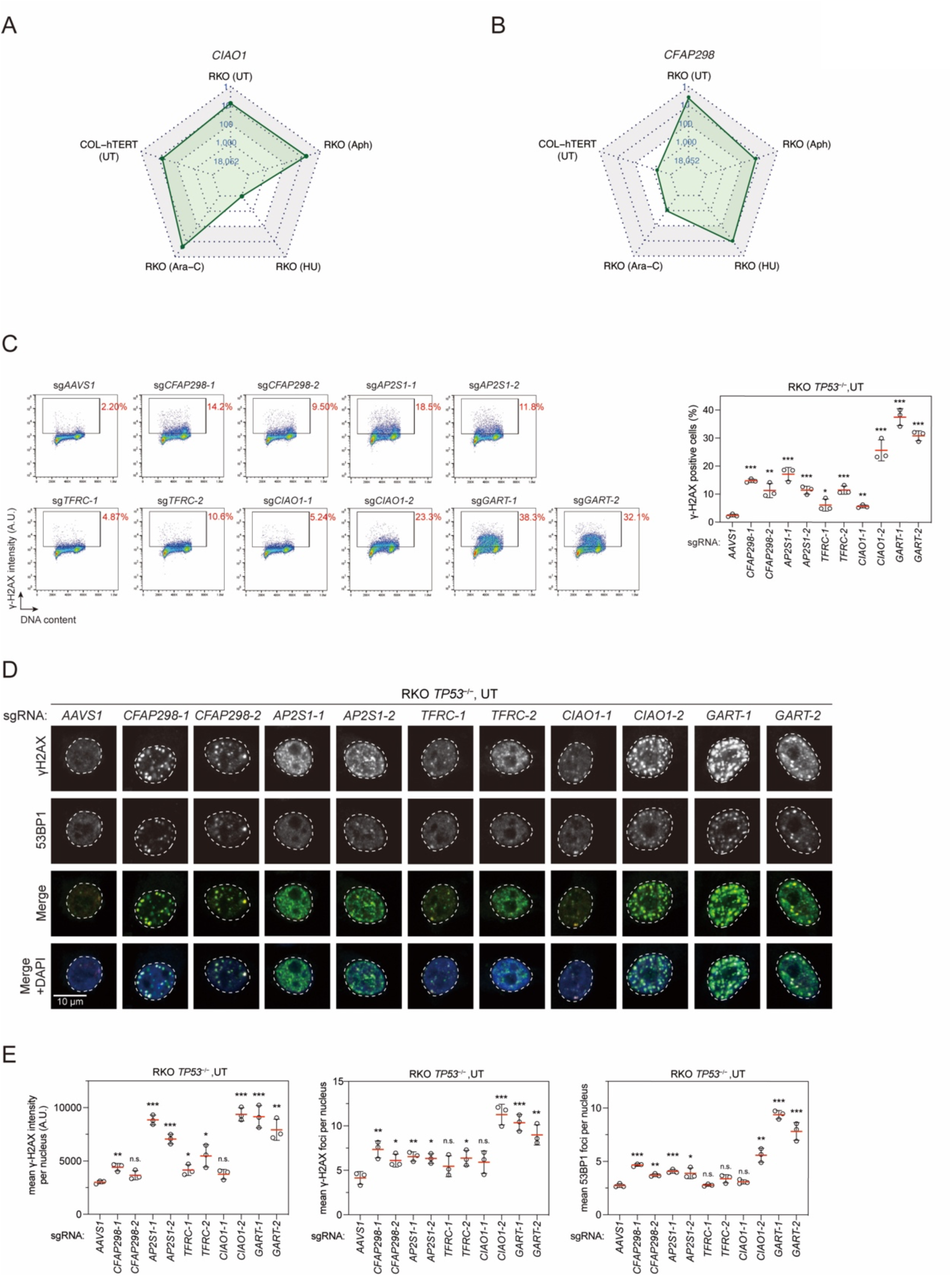
Characterization of genes that prevent endogenous DNA damage formation. (A) A radar plot showing the ranking of *CIAO1* in all five γ-H2AX screens. Custom scaling was used for five rings: 1, 10, 100, 1000, 18052, and linear scaling was used within each section. Grey shaded area indicates top 100 ranking in each screen. (B) A radar plot showing the ranking of *CFAP298* in all five γ-H2AX screens. (C) Flow cytometry analysis of RKO *TP53^−/−^* cells infected with lentiviruses expressing the indicated sgRNA. Eight days after infection, cells were fixed and stained with a γ-H2AX antibody and DAPI to monitor DNA content. Left, representative plots. Red numbers indicate the percentage of γ-H2AX positive cells. Right, quantification of the experiment shown in left. Bars represent the mean ± s.d. (n=3 independent experiments). Comparisons are made to the sg*AAVS1* control, using an unpaired t-test (*p<0.05; **p<0.01; ***p<0.001). (D) Immunofluorescence analysis of cells described in (C) with γ-H2AX and 53BP1 antibodies. The images presented are representative of three immunostainings. Dashed lines indicate the nuclear area determined by DAPI staining. (E) Quantification of mean γ-H2AX intensity (left), mean γ-H2AX focus number (middle), and mean 53BP1 focus number (right) of three independent experiments as shown in (D). Each experiment includes a minimum of 500 cells for analysis. Bars represent the mean ± s.d. Comparisons were made to the sg*AAVS1* condition using an unpaired t-test (*p<0.05; **p<0.01; ***p<0.001, n.s., not significant). A.U., arbitrary units.

To validate the results obtained in the untreated RKO *TP53*^−/−^ cell line, we selected 5 genes whose mutation increases γ-H2AX levels: the aforementioned *CIAO1* and *CFAP298* genes (Fig 2A, B), the gene encoding the AP-2 complex component AP2S1, the gene coding for the transferrin receptor TFRC involved in iron uptake (also known as CD71) and *GART*, which codes for an enzyme involved in de novo purine biosynthesis (Fig S3).

Using two independent sgRNAs, we observed that disruption of all five genes caused higher γ-H2AX signal compared to the *AAVS1-*targeting sgRNA control, confirming the screen results (Fig 2C). Examination of the γ-H2AX subnuclear localization using immunofluorescence microscopy revealed varied staining patterns for γ-H2AX and for 53BP1, a marker of DNA double-strand breaks (DSBs; Fig 2D, E). These results suggest that these genes impact genome maintenance via distinct mechanisms. For example, disruption of *CFAP298* caused mainly γ-H2AX and 53BP1 subnuclear foci, which are indicative of DSBs. In contrast, sgRNAs targeting *AP2S1* and *TFRC* induced a pan-nuclear γ-H2AX signal with little increase in γ-H2AX and 53BP1 foci, suggesting DNA replication stress rather than DSBs. Depletion of *CIAO1* and *GART* produced both pan-nuclear and focal staining of γ-H2AX, suggesting that these genes guard against both replication stress and DSB formation.

### FANCJ protects cells from Aph-induced replication stress

To identify genes that protect cells against replication stress-induced DNA damage, we searched for genes that increased γ-H2AX levels under conditions when cells were treated with DNA replication inhibitors. This analysis identified 227 genes (Supplementary Table 2) that displayed both highly selective profiles for a single DNA replication inhibitor or showed a response to multiple agents. Below we highlight a few genes along with a more in-depth characterization of the uncharacterized *GNB1L* gene.

One gene that displayed a surprisingly selective profile was *FANCJ* (also known as *BACH1* or *BRIP1*), which ranked very highly in the Aph treatment screen but not in the others (Fig 3A). FANCJ is a well-characterized DNA helicase first identified as a BRCA1-interacting protein involved in HR repair of DSBs (Cantor et al. 2001). Biallelic *FANCJ* mutations cause Fanconi anemia (FA), a rare disorder characterized by bone marrow failure and cancer predisposition (Levitus et al. 2005; Levran et al. 2005). We confirmed that cells expressing *FANCJ*-targeting sgRNAs display a large increase in γ-H2AX following Aph treatment, with a comparatively much smaller increase following Ara-C treatment (Fig 3B). Interestingly, a parallel CRISPR/Cas9 fitness screen in RKO *TP53*^−/−^ cells aimed at identifying genes whose mutation cause sensitivity to a low dose of Aph identified sgRNAs targeting *FANCJ* as the top sensitizers (Fig S4A), indicating that FANCJ has an important role in mitigating the impact of DNA polymerase inhibition in human cells.

**Figure 3.**
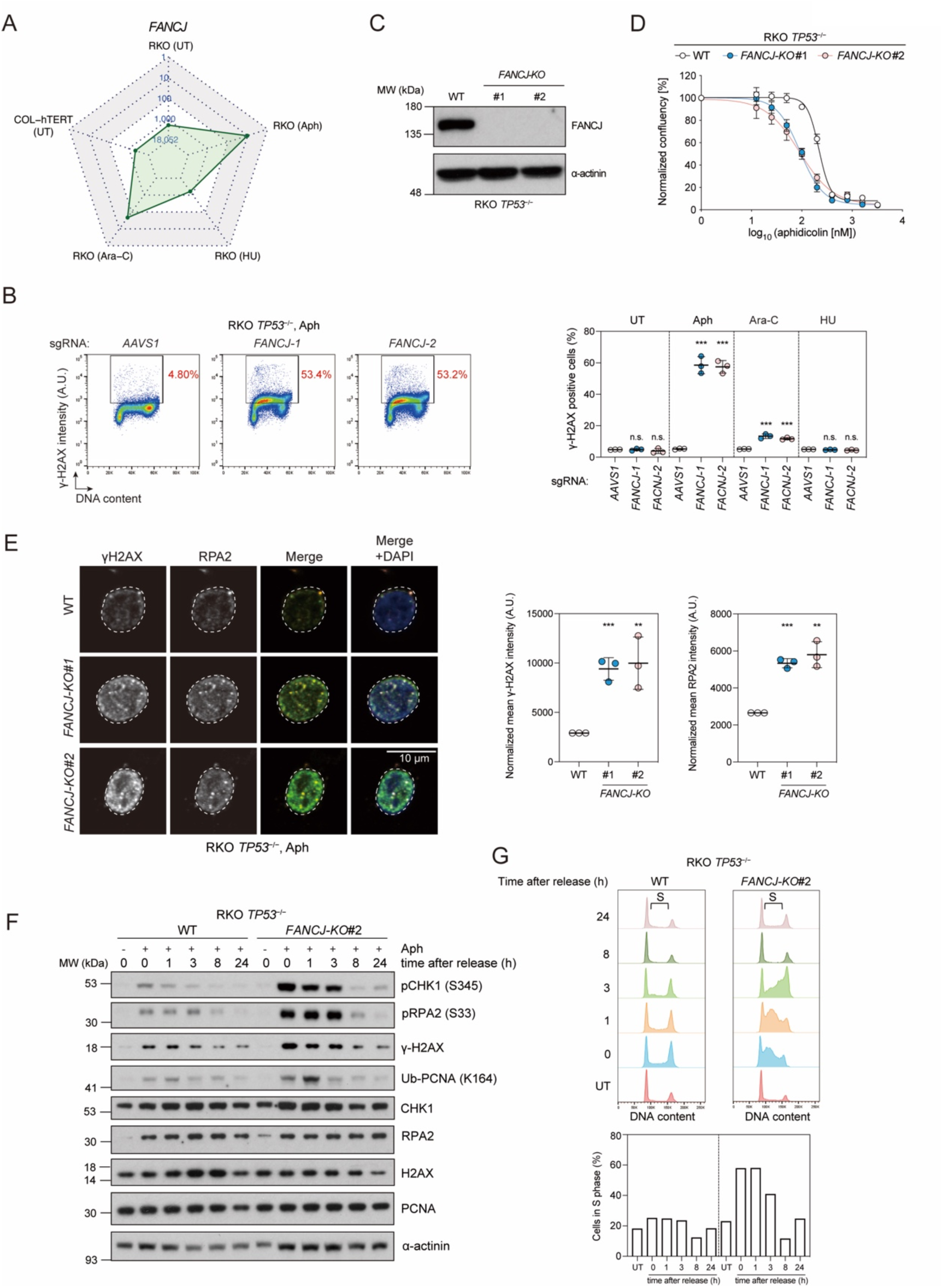
*FANCJ* protects cells from Aph-induced replication stress. (A) A radar plot showing the ranking of *FANCJ* in all five γ-H2AX screens. UT, untreated. (B) Flow cytometry analysis of RKO *TP53^−/−^* cells expressing the indicated sgRNA. Cells were treated with the indicated replication inhibitor for 24 h or left untreated (UT). 300 nM Aph, 200 nM Ara-C, and 200 μM HU were used in this experiment and the same drug concentrations were used for subsequent experiments unless otherwise specified. After fixation, cells were stained with a γ-H2AX antibody and DAPI to monitor DNA content. Left, representative flow cytometry plots following Aph treatment. Red numbers indicate the percentage of γ-H2AX positive cells. Right, quantification of γ-H2AX positive cells in all four conditions. Bars represent the mean ± s.d. (n=3 independent experiments). Comparisons are made to the sg*AAVS1* control within each treatment condition, using an unpaired t-test (***p<0.001; n.s., not significant). (C) Immunoblot analysis of FANCJ expression in RKO *TP53^−/−^* parental (WT) and *FANCJ-KO* cells. α-actinin was used as a loading control. (D) Dose-response assays with Aph in RKO WT and *FANCJ-KO* cells using confluency as a readout 6 d post-treatment. Bars represent the mean ± s.d. (n=3 independent experiments). (E) Left, immunofluorescence analysis of γ-H2AX and chromatin-bound RPA2 in RKO WT and *FANCJ-KO* cells. Cells were subjected to nuclear extraction prior to fixation. Dashed lines indicate the nuclear area determined by DAPI staining. Right, quantification of normalized mean intensities of γ-H2AX and RPA2 in three independent experiments. Each experiment includes a minimum of 500 cells for analysis. Bars represent the mean ± s.d. (n=3 independent experiments). Results of unpaired t-test between WT and *FANCJ-KO* cells are shown (**p<0.01; *** p<0.001). (F), (G) Recovery assay from Aph treatment. Cells were left untreated (−) or treated with 300 nM Aph for 24 hours, then released into cell growth medium without drug for the indicated time before harvesting. (F) Immunoblot analysis using antibodies to phospho-CHK1, phospho-RPA2, γ-H2AX, monoubiquitinated PCNA (K164), and the unmodified versions of these proteins. α-actinin, loading control. (G) Cell cycle distributions as determined by DAPI staining at the indicated times post-release from Aph treatment. UT, untreated. Brackets indicate S-phase cells. A.U., arbitrary units.

To gain insights into the replication defects of *FANCJ*-deficient cells, we generated CRISPR knockout (KO) clones of *FANCJ* in RKO *TP53*^−/−^ cells, which were confirmed by immunoblot analysis (Fig 3C). *FANCJ-KO* cells are hyper-sensitive to Aph, as expected (Fig 3D). Under Aph challenge, *FANCJ-KO* cells accumulate in S phase (Fig S4B) and show a striking increase in pan-nuclear γ-H2AX staining accompanied by concomitant increase in chromatin-bound RPA2, which is part of the ssDNA-binding complex RPA (Fig 3E). These results suggest that FANCJ loss leads to widespread replication perturbation characterized by generation of ssDNA.

The formation of ssDNA in *FANCJ-KO* cells following a 24 h Aph treatment was accompanied with increased ATR signaling, as monitored through CHK1-S345 and RPA2-S33 phosphorylation (Fig 3F). Following termination of the Aph treatment, ATR signaling gradually decreased to reach baseline levels 8 h post-release. Similarly, the striking S-phase accumulation of *FANCJ-KO* cells in the presence of low-dose Aph was resolved within 8 h, with cell cycle profiles becoming undistinguishable from parental cells (Fig 3G). These observations suggest that the DNA lesions in *FANCJ-KO* cells are largely reversible.

In budding yeast, ssDNA gaps can be repaired following the recruitment of translesion synthesis (TLS) polymerases (Gallo et al. 2019), which are recruited to DNA lesions by PCNA ubiquitylation (Choe and Moldovan 2017). We monitored monoubiquitylation of PCNA Lys164 after Aph treatment and release and observed that *FANCJ-KO* cells displayed increased PCNA ubiquitylation that rapidly peaked 1 h post-release (Fig 3F). Together, these observations suggest that under Aph challenge, *FANCJ-*deficient cells accumulate ssDNA gaps that are likely reversed in part by the action of TLS polymerases.

The γ-H2AX accumulation displayed by *FANCJ-KO* cells during Aph treatment can readily be rescued by lentiviral expression of wild-type FANCJ (Fig S4C-E). This allowed us to profile various FANCJ mutants to gain insights into the mechanism by which FANCJ suppresses ssDNA during low-dose Aph treatment. We introduced into *FANCJ-KO* cells the following FACNJ mutants: K52R, which disrupts helicase activity (Cantor et al. 2001); K141A,K142A which impairs interaction with MLH1 (Peng et al. 2007); S990A, which abolishes interaction with BRCA1 (Manke et al. 2003); and T1133A, which blocks binding to TOPBP1 (Gong et al. 2010). Reintroduction of these mutants, with the notable exception of FANCJ-K52R, suppressed the γ-H2AX accumulation in the presence of Aph to the levels observed with wild-type FANCJ (Fig S4C-E). Identical results were obtained in RPE1-hTERT cells where the FANCJ mutations were introduced at the chromosomal locus via gene editing (Fig S4F). These results suggest that the FANCJ helicase activity, but not its roles in ICL repair, HR or DNA damage signaling, is essential to protect cells from DNA replication stress.

### DERA and PGD act against Ara-C-induced replication stress

*DERA* and *PGD* were the top-ranking genes in the Ara-C γ-H2AX screen (Fig S5A, B) and we validated that mutation of either gene causes high levels of γ-H2AX following Ara-C treatment (Fig S5C, D). DERA is the human deoxyribose phosphate aldolase, which participates in various metabolic pathways including nucleotide catabolism and the pentose phosphate pathway (Salleron et al. 2014). It catalyzes the conversion of 2-deoxy-D-ribose-5-phosphate into glyceraldehyde-3-phosphate and acetaldehyde (Salleron et al. 2014). *PGD* encodes the enzyme 6-phosphogluconate dehydrogenase, the third enzyme in oxidative pentose phosphate pathway that converts 6-phosphogluconate to ribulose 5-phosphate and produces NADPH (Lin et al. 2015). DERA is specifically required to prevent DNA damage induced by Ara-C, but not by Aph or HU (Fig S5C). *DERA-*targeting sgRNAs greatly impair cell proliferation in the presence of low dose Ara-C but not in untreated cells (Fig S5E). Given the role of DERA in nucleotide salvage pathways, it may play a previously unsuspected role in the detoxification of Ara-C in human cells.

### RECQL5 protects cells from DNA double-strand breaks under replication stress

*RECQL5* is representative of a gene whose mutation increased γ-H2AX levels selectively under the three types of DNA replication stress tested, with a phenotype that was stronger following Aph treatment (Fig S6A). *RECQL5* encodes a RecQ-type 3’-5’ DNA helicase (Fig S6B) implicated in DNA repair and replication (Garcia et al. 2004). It associates with the replisome in S phase and persists at sites of stalled replication forks (Kanagaraj et al. 2006). Independent sgRNAs targeting *RECQL5* caused increased γ-H2AX in response to Aph, HU or Ara-C when compared to controls, confirming the screen results (Fig S6C). RECQL5-deficient cells accumulated γ-H2AX and 53BP1 foci following Aph treatment, suggesting the formation of DSBs (Fig S6D). A clonal knockout of *RECQL5* generated by gene editing (Fig S6E,F) also showed elevated γ-H2AX levels following Aph treatment, a phenotype that we could suppress by reintroduction of *RECQL5* with lentiviral transduction (Fig S6E,F). This system allowed us to test a number of characterized RECQL5 variants that disrupt either its helicase activity (K58R), its phosphorylation by CDK1 (S727A) or its interaction with RAD51 (F666A) or RNA polymerase II (E584D) (Garcia et al. 2004; Schwendener et al. 2010; Di Marco et al. 2017). To our surprise, every RECQL5 mutant complemented *RECQL5-KO* cells to the same extent as wild type RECQL5 (Fig S6B, E, F). These results suggest that the mechanism by which RECQL5 prevents DSB formation following replication stress may depend on an activity that is currently uncharacterized.

### GNB1L prevents replication catastrophe under mild replication stress

*GNBL1* is another gene whose mutation caused high γ-H2AX levels following treatment with replication inhibitors, with a particularly strong response in the HU screen (Fig 4A). *GNB1L* was of particular interest since it has been repeatedly linked to schizophrenia and autism in genetic association studies (Williams et al. 2008; Ishiguro et al. 2010; Chen et al. 2012), and *Gnb1l* knockout mice are embryonic lethal (Paylor et al. 2006). *GNB1L* encodes a protein of unknown function consisting of seven predicted WD40 repeats (Gong et al. 2000; Funke et al. 2001).

**Figure 4.**
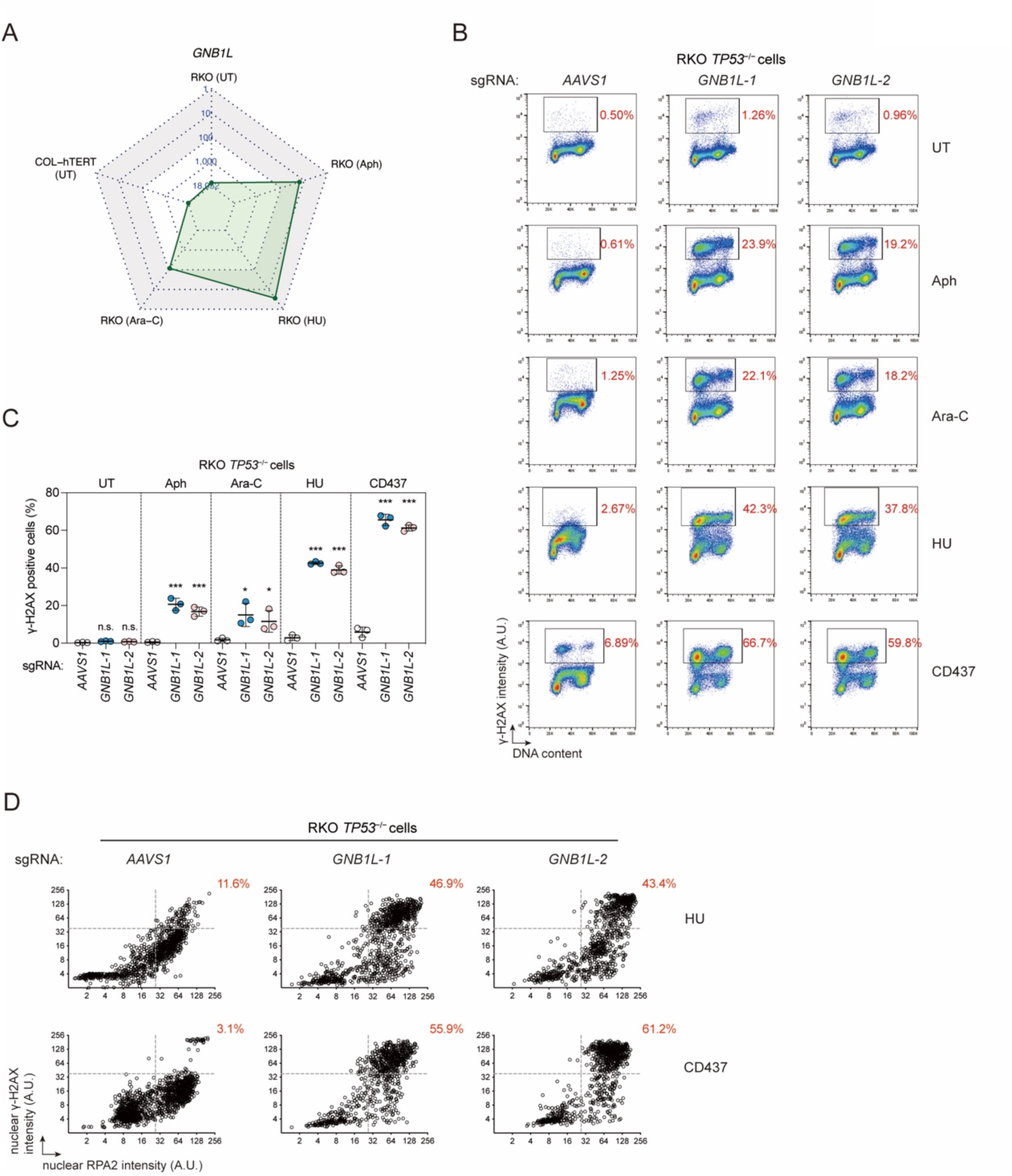
*GNB1L* protects RKO *TP53^−/−^* cells from replication catastrophe under mild replication stress. (A) A radar plot showing the ranking of *GNB1L* in all five γ-H2AX screens. (B) Representative flow cytometry analysis of RKO *TP53^−/−^* cells expressing the indicated sgRNA. Cells were treated with the indicated replication inhibitor for 24 h or left untreated (UT), and then fixed, stained with a γ-H2AX antibody and DAPI. Red numbers indicate the percentage of γ-H2AX positive cells. (C) Quantification of the experiment shown in (B). Bars represent the mean ± s.d. (n=3 independent experiments). Comparisons are made to the sg*AAVS1* control within each treatment condition, using an unpaired t-test (***p<0.001; n.s., not significant). (D) Quantitative image-based cytometry (QIBC) analysis of γ-H2AX and chromatin-bound RPA2 signal intensities in RKO *TP53^−/−^* cells. Cells were treated with 200 μM HU or 250 nM CD437 for 24 hours, then extracted, fixed and stained with antibodies to γ-H2AX and RPA2. Red numbers indicate the percentage of cells with both high γ-H2AX and high RPA2 signal intensities for each condition. A.U., arbitrary units.

RKO *TP53^−/−^* and RPE1-hTERT *TP53^−/−^* cells expressing *GNB1L-*targeting sgRNAs display a massive increase in γ-H2AX in S-phase cells, specifically under conditions of mild replication stress (Figs 4B,C and S7A). The induction of γ-H2AX following HU treatment was reminiscent of replication catastrophe, a condition caused by extensive ssDNA triggers exhaustion of the RPA pool, leading to unprotected ssDNA and subsequent DSB formation (Toledo et al. 2013). To explore whether GNB1L guards against replication catastrophe, we monitored the response to the DNA polymerase α inhibitor CD437 (Han et al. 2016), which is particularly efficient at eliciting replication catastrophe (Ercilla et al. 2020). GNB1L loss greatly potentiated γ-H2AX induction under low doses of CD437 (Figs 4B,C and S7A). To formally assign the phenotype to replication catastrophe, we performed quantitative image-based cytometry (QIBC) to simultaneously monitor H2AX phosphorylation and the extent of ssDNA formation by measuring the levels of chromatin-bound RPA. *GNB1L* depletion caused the accumulation of cells that displayed both high γ-H2AX and chromatin-bound RPA signals, characteristic of replication catastrophe (Figs 4D and S7B). GNB1L therefore guards against DNA replication catastrophe.

### GNB1L interacts with PIKKs and the TTT complex

To gain insights into the mechanism by which GNB1L protects cells from replication stress, we searched for interacting proteins using two parallel approaches. We employed affinity purification coupled to mass spectrometry (AP-MS) and proximity-based interaction proteomics using the BirA-related miniTurbo system (Branon et al. 2018). After filtering hits with SAINT (Teo et al. 2014) we found that TELO2 was the only high-confident protein interacting with GNB1L in the AP-MS experiment (Supplementary Table 4). In contrast, proximity interaction proteomics also retrieved TELO2 but did so alongside TTI1 and all six members of the PIKK family (DNA-PKcs, ATM, ATR, mTOR, SMG1 and TRRAP) (Fig 5A and Supplementary Table 4). Streptavidin pulldowns followed by immunoblot analysis in cells expressing miniTurbo-tagged GNB1L (Fig S8A) confirmed that each the TTT complex and each PIKK protein resided in the proximity of GNB1L. Similarly, the same proteins, with the exception of SMG1, could be retrieved in 3xFlag-GNB1L immunoprecipitates, suggesting more intimate interactions than anticipated from their AP-MS profiles (Fig 5B). Finally, we developed a NanoBRET assay (Machleidt et al. 2015) that further confirmed a physical interaction between GNB1L and TELO2, suggesting that these two proteins are closely associated (Fig S8B).

**Figure 5.**
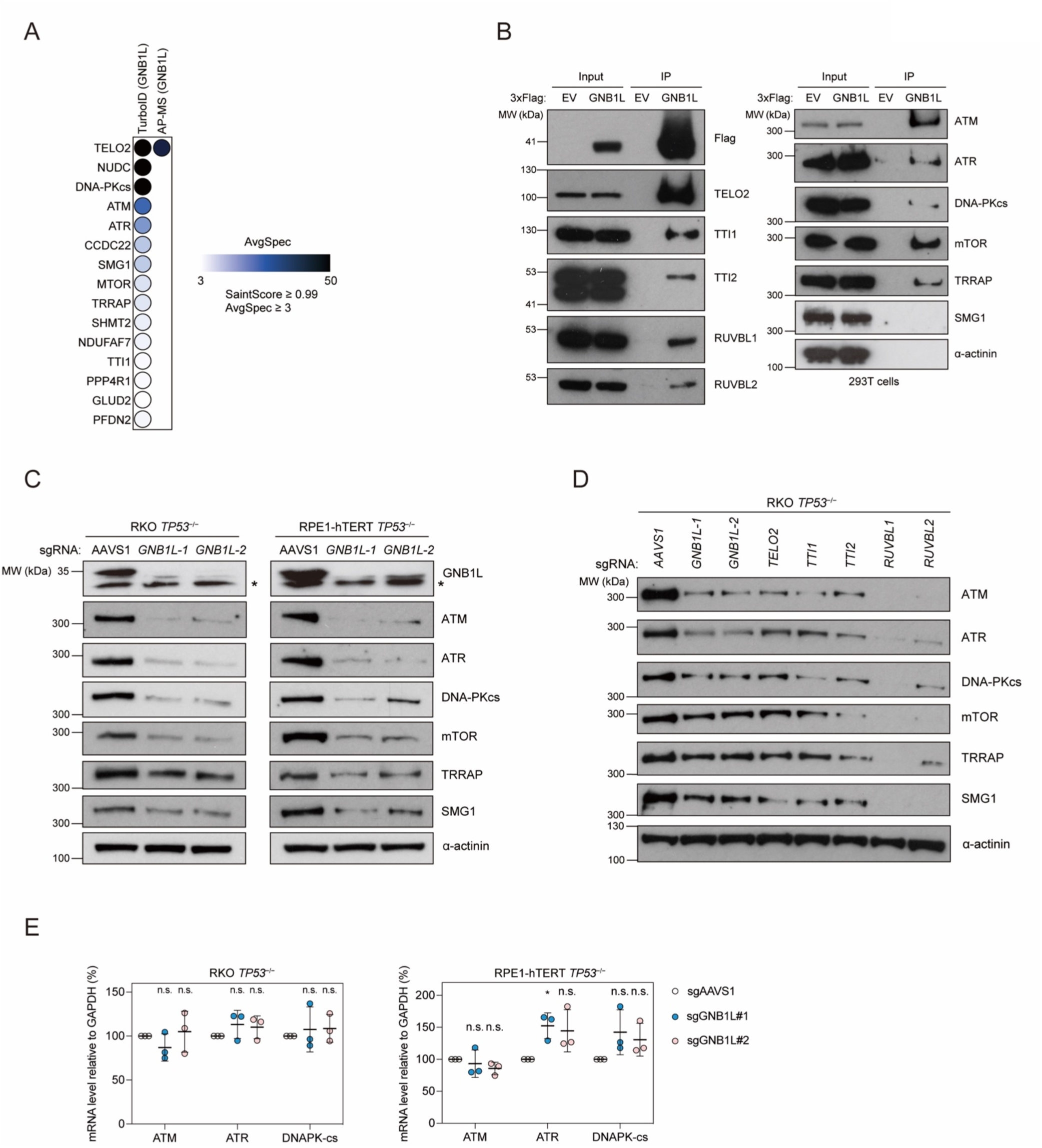
GNB1L physically interacts with PIKKs and is essential to maintain their protein levels. (A) Mass spectrometry results of the TurboID-based proximal labeling experiment with GNB1L-miniTurbo and AP-MS with 3xFlag-GNB1L experiments. High-confident hits with a SaintScore ≥ 0.99 and average spectral counts ≥ 3 are shown for both experiments. Color indicates the number of average spectral counts minus the control counts. (B) Anti-Flag immunoprecipitation in 293T cells expressing 3xFlag-GNB1L or 3xFlag alone (EV). Bound proteins were examined by immunoblot analysis with the indicated antibodies. (C) Immunoblot analysis of PIKK proteins in lysates from RKO *TP53^−/−^* and RPE-hTERT *TP53^−/−^* cells expressing sg*AAVS1* control or sg*GNB1L*. α-actinin, loading control. Asterisk besides GNB1L immunoblots indicate non-specific bands. (D) Immunoblot analysis of PIKK proteins in lysates from RKO *TP53^−/−^* cells expressing the indicated sgRNA. α-actinin, loading control. (E) Quantitative RT-PCR to detect mRNA of ATM, ATR, DNA-PKcs using TaqMan assays. Bars represent the mean ± s.d. (n=3 independent experiments). Results of an unpaired t-test between sg*AAVS1* and sg*GNB1L* conditions are shown (*p<0.05).

### GNB1L promotes PIKK protein biogenesis

The identification of TTT, RUVBL1/2 and PIKKs as GNB1L-interacting proteins was revealing because the replication catastrophe seen in GNB1L-deficient cells is a phenocopy of the phenotype caused by ATR loss or inhibition (Toledo et al. 2013). Given that TTT and RUVBL1/2 promote ATR (and PIKK) biogenesis, these results suggested that GNB1L may also participate in the biogenesis of PIKKs. Indeed, transduction of *GNB1L*-targeting sgRNAs caused a steady-state reduction in the levels of all PIKKs, with the DNA damage-related factors, ATM, ATR and DNA-PKcs being the most affected (Fig 5C). The reduction in PIKK levels was similar to that observed in cells expressing sgRNAs targeting genes encoding members of the TTT complex (Figs 5D and S8C), which also caused replication catastrophe-like phenotype under mild replication stress (Fig S8D). The reduced PIKK levels in GNB1L-depleted cells was not due to reduced mRNA levels, at least for the 3 PIKKs tested (Fig 5E). To assess if the reduction in PIKK protein levels was translated into impaired function, we assessed the integrity of ATM, ATR and mTOR signaling by immunoblot analysis. Substrate phosphorylation by ATR, ATM, and mTOR was compromised in *GNB1L*-depleted cells after their stimulation (Fig S8E). Finally, consistent with ATR signaling being critical for the viability of cells experiencing DNA replication stress (Saldivar et al. 2017), we observed that *GNB1L*-deficient cells displayed impaired proliferation in the presence of HU or Ara-C (Fig S8F). Together, these results indicate that GNB1L promotes the biogenesis of functional PIKK proteins.

TELO2 stabilizes newly synthesized but not pre-existing PIKKs (Takai et al. 2010). To test whether GNB1L also selectively promotes the biogenesis of newly synthesized PIKKs, we first generated a 293T cell line derivative in which the GNB1L protein can be rapidly depleted using the dTAG system (Nabet et al. 2018; Nabet et al. 2020) (Fig S9A). Over time, GNB1L depletion caused a reduction in the protein levels of ATM, ATR, DNA-PKcs, as well as sensitizing cells to replication catastrophe (Fig S9B). To monitor ATR biogenesis in this cell line, we expressed Halo-tagged ATR, which enabled us to follow the abundance of newly synthesized or pre-existing ATR using label-switch strategies (Fig S9C, D). With this system, we found that GNB1L depletion selectively impacted the accumulation of newly synthesized ATR (Fig S9C, D). We conclude that GNB1L, like TELO2, promotes the biogenesis of newly synthesized PIKKs, including that of ATR.

### Deep mutational scanning of GNB1L

The observation that GNB1L is predicted to form a single WD40 repeat propeller, which represents a folded unit, suggested that deletion mutagenesis as a means to identify functionally important regions of GNB1L would be impossible. As an alternative, we applied deep mutational scanning to identify variants of GNB1L that promote replication catastrophe (Fowler and Fields 2014). We constructed a lentiviral *GNB1L* mutant library of 5529 mutants by POPcode mutagenesis (Weile et al. 2017) in which GNB1L is expressed as a C-terminal GFP fusion (Fig 6A). After library transduction, we sorted for GFP-positive cells to remove nonsense variants or variants that caused loss of GNB1L expression. Endogenous *GNB1L* was then inactivated with an sgRNA targeting an intron-exon junction. The resulting pool of cells was treated with HU and subsequently sorted for cells with high γ-H2AX (Fig. 6A). Mutant frequency in both the unsorted cell populations was determined using Tileseq (Weile et al. 2017; Sun et al. 2020). A “functional” score was assigned to each mutant (Supplementary Table 5) based on their relative abundance in the high γ-H2AX cell population; mutations with low functional scores suggest that they compromise the ability of GNB1L to prevent replication catastrophe.

**Figure 6.**
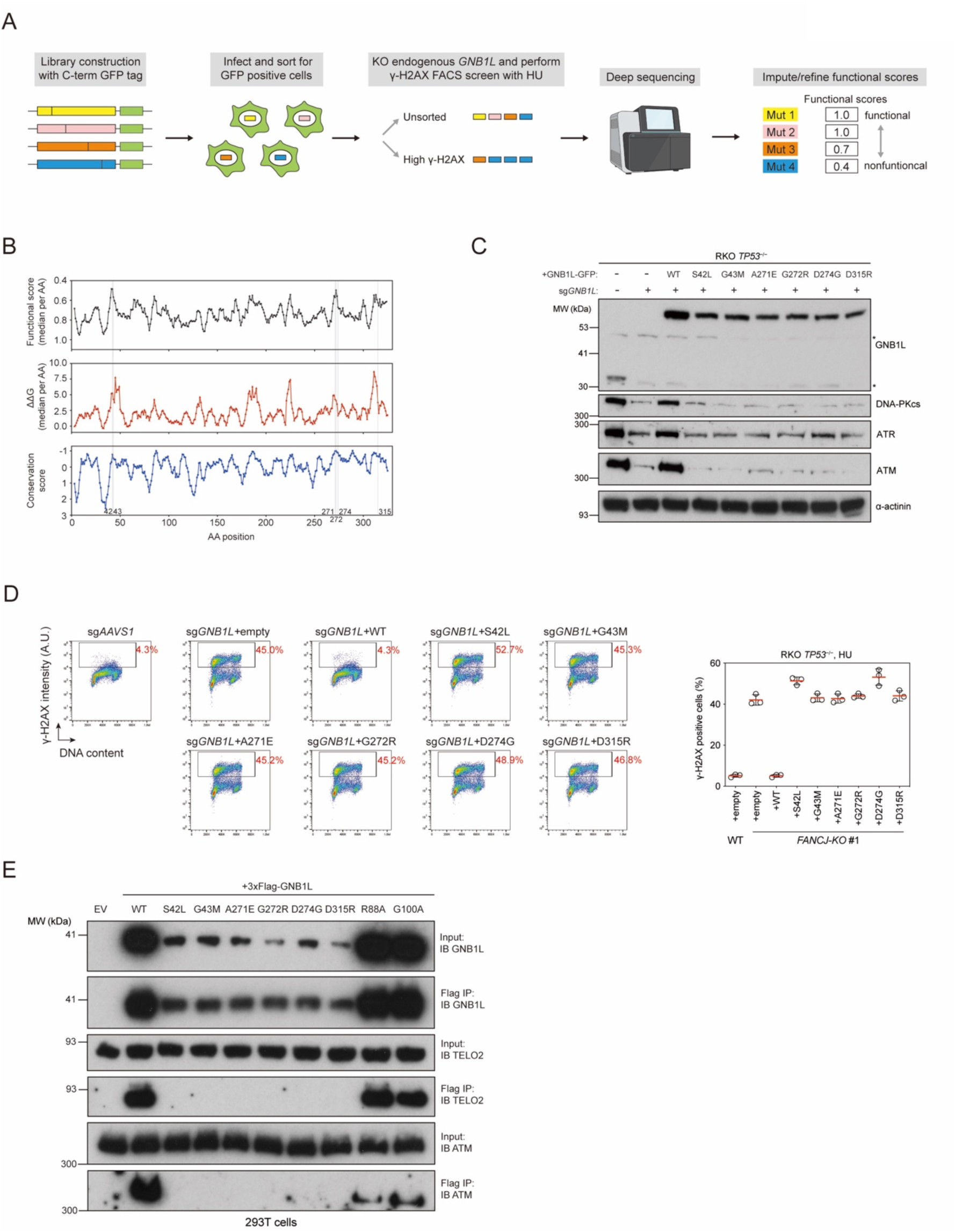
Deep mutational scanning of the GNB1L protein. (A) Schematic of the deep mutational scanning strategy. (B) The moving window analysis of the median functional score, median ΔΔG, and the conservation score for each residue of GNB1L, with a window of five residues. (C), (D) RKO *TP53^−/−^* cells were infected with lentiviruses expressing sg*GNB1L* and sgRNA-resistant GNB1L-GFP variant constructs as indicated. WT, wildtype. (C) Immunoblot analysis of cell lysates with the indicated antibodies. α-actinin, loading control. (D) Cells were treated with 250 nM CD437 for 24 hours, then fixed and stained with a γ-H2AX antibody and DAPI. Left, representative flow cytometry plots. Right, quantification of γ-H2AX positive cells. Bars represent the mean ± s.d. (n=3 independent experiments). A.U., arbitrary units. (E) Anti-Flag immunoprecipitation (IP) in 293T cells expressing 3xFlag-tagged GNB1L (WT or mutants). WT, wildtype. EV, empty vector. Bound proteins were examined by immunoblotting (IB) with the indicated antibodies.

In deep mutation scanning studies such as variant effect maps, nonsense mutations are useful landmarks for calibrating variant effects and for quality control. However, as we selected against nonsense variants in the GFP sorting step, we examined the relationship between the median functional score for each GNB1L residue and a corresponding median change in Gibb’s free energy (ΔΔG) computed with FoldX (Guerois et al. 2002) (Fig 6B and Supplementary Table 6). Using a moving window analysis, we observed an inverse correlation between functional scores and the median ΔΔG (Pearson correlation −0.61) suggesting that substitutions that perturb folding are functionally impaired, as expected. Similarly, using conservation computed with ConSurf (Fig 6B and Supplementary Table 6), where negative values indicate conservation, we observed positive correlation between functional and conservation scores (Pearson correlation 0.58). Together, these analyses indicate that our mutational scanning pipeline is effective at identifying variants that impair GNB1L function.

We next mined the deep mutational scanning dataset to identify variants that have low ΔΔG scores (i.e. have minimal impact on protein folding) but have large impact on GNB1L function (low functional scores). We selected 6 high-confidence variants for retesting: S42L, G43M, A271E, G272R, D274G and D315R (Supplementary Table 5). Unlike wild type GNB1L, which can rescue replication catastrophe caused by a *GNB1L*-targeting sgRNA, reintroduction of each of the variants failed to rescue HU-induced γ-H2AX levels, indicating that they were all defective in preventing replication catastrophe (Fig 6C, D). While the expression of the GNB1L variants was lower than that of exogenously expressed GNB1L, they were all expressed at higher levels than endogenous GNB1L in the parental cell line (Fig 6C), displayed lower levels of ATM, ATR and DNA-PKcs proteins (Fig 6C) and all were impaired in their interaction with TELO2 (Fig 6E) suggesting that the integrity of the GNB1L-TELO2 complex is critical for PIKK biogenesis. Exactly how these mutations impact GNB1L function is not clear but as they are not concentrated on any one area of the proteins, it is likely that at least a subset of them act by subtly destabilizing the GNB1L structure.

### The GNB1L-TELO2 interaction is essential for PIKK stability

In parallel, we mapped the region of TELO2 involved in its interaction with GNB1L. The region of TELO2 that is necessary and sufficient to interact with GNB1L is encompassed by residues 460-545 (Figs 7A and S10A, B), which is consistent with an AlphaFold model of the GNB1L-TELO2 complex that identified a TELO2 loop encompassing these residues as directly interacting with GNB1L (Jumper et al. 2021) (Fig 7B, S10C, D, and Supplementary Table 7). Alanine scanning of this region identified a TELO2 mutant we designated as M2 (^498^YMDS^501^-AAAA) that produced a variant expressed at wild-type levels that was completely deficient in GNB1L binding, and in protecting cells against replication catastrophe, and which displayed impaired PIKK levels (Fig 7C, D). This was in contrast to single point mutants of ^498^YDMS^501^, which were impaired in TELO2 binding but displayed both normal PIKK levels and responses to HU treatment (Fig S10E-G). These results indicate that a minimal TELO2-GNB1L interaction is both sufficient and necessary for PIKK stability and guards against replication catastrophe.

**Figure 7.**
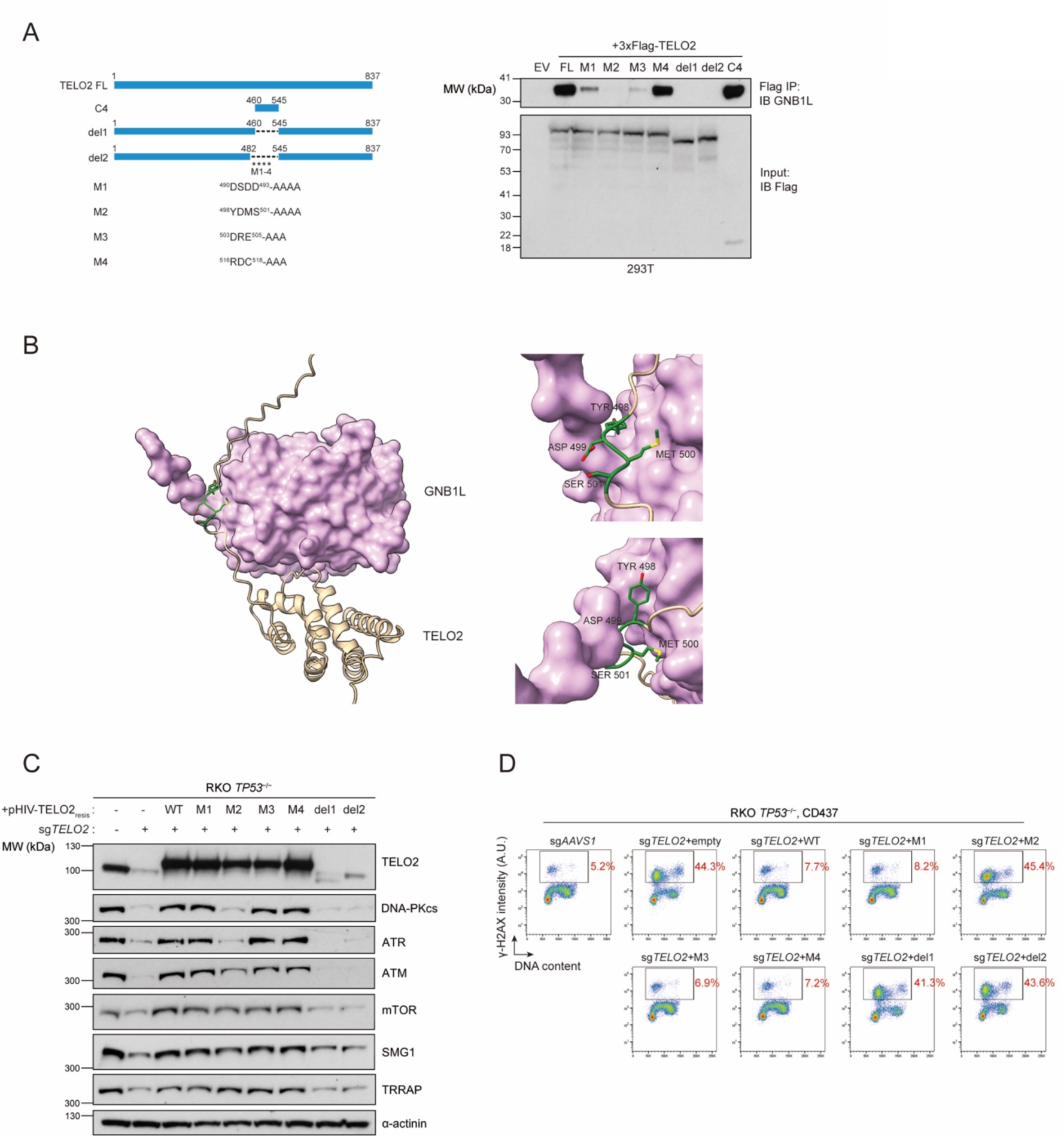
GNB1L functions with TTT complex for PIKK biosynthesis. (A) Left, Schematic of truncation/point mutation mapping in the TELO2 protein. Right, Anti-Flag immunoprecipitation (IP) in 293T cells expressing full length (FL) or mutant 3xFlag-TELO2. The GNB1L-TELO2 interaction was examined by immunoblotting with a GNB1L antibody. (B) Left, AlphaFold2-predicted structure of full-length GNB1L binding to the C3 fragment of TELO2 (460-640aa). Purple, surface structure of GNB1L. Beige, ribbon structure of the TELO2 fragment. Right, magnified view of binding surface. Amino acid residues 498-501 of TELO2 are labeled and highlighted in green. (C), (D) RKO *TP53^−/−^* cells were infected with lentiviruses expressing sg*TELO2* and sgRNA-resistant TELO2 variant constructs as indicated. WT, wildtype. (C) Immunoblot analysis of cell lysates with the indicated antibodies. α-actinin, loading control. (D) Cells were treated with 250 nM CD437 for 24 hours, and then fixed, stained with a γ-H2AX antibody and DAPI. Red numbers indicate the percentage of γ-H2AX positive cells. The results are representative of two independent experiments. A.U., arbitrary units.

## Discussion

This work presents a genome-scale survey of the genes and pathways that protect cells against DNA damage using H2AX phosphorylation as a readout. Our analysis identified 160 genes that protect cells from spontaneous DNA lesions in RKO *TP53^−/−^* cells, with an additional 227 genes that suppress DNA damage under conditions of mild DNA replication stress. This dataset can be mined for the identification of proteins not previously associated with genome maintenance. For example, we identified *CFAP298*, a gene previously linked to ciliogenesis (Austin-Tse et al. 2013), as suppressing the formation of spontaneous DNA double-strand breaks.

One of the most surprising findings of this study was that the 160 genes suppressing DNA damage were functionally enriched in only handful of biological processes. Those include DNA replication, DNA repair, RNA metabolism and a few biosynthetic pathways such as nucleotide metabolism and Fe-S cluster biogenesis. These results suggest that the genome may be insulated from the majority of cellular processes, and we speculate that this could effectively minimize the impact of dysfunctional cellular processes on the integrity of the genome.

Among the genes that increased γ-H2AX levels in RKO *TP53^−/−^* cells under unchallenged conditions, 14 genes encode proteins participating in Fe-S cluster assembly and three genes (*AP2S1*, *TFRC*, *FCHO2*) code for factors involved in iron uptake by endocytosis. This observation further highlights a key role for iron metabolism in genome stability, which can be explained by the fact that many DNA replication and repair proteins require a Fe-S cluster as their cofactor, as previously noted (Paul and Lill 2015) and by the fact that the diferric-tyrosyl radical central to ribonucleotide reductase (RNR) activity also depends on the Fe-S cluster assembly pathway for its maturation (Li et al. 2017).

Although mitochondrial dysfunction is often assumed to cause nuclear DNA damage as a consequence of oxidative stress (Babbar et al. 2020), our results point to defective Fe-S cluster assembly as an important source of nuclear genome damage. This is consistent with work in budding yeast that linked the genome instability caused by age-related mitochondrial dysfunction to defective Fe-S cluster assembly (Veatch et al. 2009). These observations also suggest that modulating iron uptake or Fe-S cluster assembly could be used to induce DNA damage for therapeutic purposes. In line with this possibility, we recently identified the mitochondrial iron transporter SLC25A28 (mitoferrin-2) as essential in *BRCA1/BRCA2*-deficient cells (Adam et al. 2021). A challenge will be, however, to design strategies that minimize the impact on physiological processes that require iron uptake, such as erythropoiesis.

We uncovered a role for the FANCJ and RECQL5 proteins in suppressing the formation of γ-H2AX-associated DNA lesions in the presence of mild DNA replications stress. While both proteins are well-characterized DNA repair factors, our structure-function studies indicate that their role in suppressing replication-associated DNA damage may be distinct from their better described function. For example, FANCJ participates in interstrand crosslink repair in a manner that requires is interaction with MLH1 (Peng et al. 2007). However, the FANCJ-MLH1 interaction is completely dispensable to suppress ssDNA accumulation in response to DNA polymerase inhibition, indicating a distinct role for FANCJ in this process. As the motor/helicase activity of FANCJ is necessary to suppress ssDNA accumulation in response to Aph treatment, our results are instead consistent with a model where FANCJ remodels DNA structures that hinder replication fork progression.

We describe GNB1L as a PIKK biogenesis factor that cooperates with the TTT-RUVBL1/2 co-chaperone complex. Since *GNB1L* haploinsufficiency is a candidate causal gene for the neuropsychiatric disorders associated with the 22q11.2 deletion syndrome (Gong et al. 2000; Paylor et al. 2006; Williams et al. 2008; Ishiguro et al. 2010; Chen et al. 2012), it is likely that defective PIKK biogenesis contributes to the pathophysiology of this syndrome. In support of this possibility, mutations in *TELO2* and *TTI2* also cause intellectual disability disorders (You et al. 2016; Ziegler et al. 2019). Exactly which PIKK or combination of PIKKs are involved the etiology of 22q11.2 syndrome (or in the *TELO2/TTI2*-associated syndromes) is unclear, but it will be useful in the future to assess PIKK signaling in *GNB1L* heterozygote mutants as a means to identify those that are sensitive to *GNB1L* gene dosage.

In budding yeast, the Tel2-Tti1-Tti2 complex promotes the protein stability of the homologs of ATR and ATM (Mec1 and Tel1, respectively) through an Asa1-dependent pathway (Goto et al. 2017). Asa1 is a WD40 repeat protein with limited homology to GNB1L, but, as noted previously, (Stirling et al. 2011) it is likely that GNB1L represents a distant Asa1 homolog in vertebrates given our described function of GNB1L. The recent elucidation of the structure of the TTT complex (Pal et al. 2021) also paves the way for studying the contribution of GNB1L to PIKK biogenesis at the biochemical and structural levels.

In addition to GNB1L, TELO2 also interacts with the PIH1D1 and RPAP3 proteins, which have also been implicated in PIKK biogenesis (Horejsi et al. 2010). PIH1D1 binds to the same region of TELO2 that associates with GNB1L but mutations in the key phosphoacceptor residues on TELO2 required for the PIH1D1 interaction (S487/S491) did not affect GNB1L binding, indicating that GNB1L forms a complex with TELO2 that does not require PIH1D1 (Fig S10E). Similarly, depletion of PIH1D1 and RPAP3 did not cause a marked decrease in PIKK protein levels in RKO *TP53^−/−^* cells (Fig S11), suggesting that the GNB1L-TTT-RUVBL1/2 pathway may be the dominant PIKK biogenesis route in human cells.

Finally, we note that catalytic inhibition of the DNA damage-associated PIKKs is clearly not equivalent to the loss of the enzymes (Menolfi and Zha 2020). For example, complete loss of ATM or DNA-PKcs is viable in mice but their kinase-dead alleles die in utero (Yamamoto et al. 2012; Jiang et al. 2015). Given that PIKK inhibition is being aggressively pursued as a therapeutic modality, it is conceivable that a strategy aimed at destabilizing PIKKs rather than their catalytic inhibition would produce distinct therapeutic and toxicity profiles. Compounds that inhibit the TELO2-GNB1L interaction or that cause GNB1L degradation may therefore be of potential interest.

## Materials and Methods

### Cell culture

RPE1-hTERT and 293T cells were obtained from ATCC. COL-hTERT cells (T0570) were purchased from ABM (Richmond, BC). 293 Flp-In cells were obtained from Invitrogen. RKO *TP53^−/−^* cells were a gift fro Agnel Sfeir at the Sloan Kettering Institute. RKO *TP53^−/−^*, RPE1-hTERT *TP53^−/−^*, COL-hTERT *TP53^−/−^*, 293T, and 293 Flp-In cells were grown in Dulbecco’s Modified Eagle Medium (DMEM; Gibco/Thermo Fisher) supplemented with 10% fetal bovine serum (FBS; Wisent), 1x non-essential amino acids, 200 mM GlutaMAX (both Gibco/Thermo Fisher), 100 U/ml penicillin and 100 μg/ml streptomycin (Pen/Strep; Wisent). All cell lines were routinely authenticated by STR and tested negative for mycoplasma.

RKO *TP53^−/−^ FANCJ-KO* and *RECQL5-KO* gene knockouts were generated by electroporation of Cas9 and sgRNA using a Lonza Amaxa II nucleofector. sgRNA target sequences were: *FANCJ*, AGATTACTAGAGAGCTCCGG; *RECQL5*, AGTCAGCTTCCTGATCAGGA. Cells were cultured for an additional five days after electroporation to provide time for gene editing and then seeded at low densities (500 cells/15-cm dish) for single-clone isolation. *FANCJ-KO* and *RECQL5-KO* cell clones were identified by PCR amplification and ICE analysis (https://ice.synthego.com) and confirmed by immunoblot analysis (Supplementary Table 8).

For the RPE1-hTERT *TP53^−/−^ FANCJ* knock-in cell lines, the desired *FANCJ* gene variants (K52R, K141/142A, S990A, T1133A) were introduced in the RPE1-hTERT *TP53^−/−^* Cas9-expressing clone, using the RNP CRISPR approach of IDT. Sequences of PCR primers, sgRNA, and ssODN repair templates can be found in Supplementary Table 9. The expression of FANCJ variants were confirmed by immunoblot analysis.

### Plasmids and viral vectors

DNA corresponding to sgRNAs was cloned into LentiCRISPRv2 using BamHI (Addgene, #52961). sgRNA target sequences and their validations can be found in Supplementary Table 8. Lentiviral particles were produced in 293T cells by co-transfection of the targeting vector with vectors expressing VSV-G, RRE and REV using calcium phosphate. Medium was refreshed 12-16 h later. Virus-containing supernatant was collected 36-40 h post transfection, cleared through a 0.4-μm filter, supplemented with 8 μg/ml polybrene (Sigma) and used for infection of target cells. pHIV-NAT-T2A-hCD52 was a kind gift of R. Scully. pcDNA5-FRT/TO-3xFLAG (V4978) was obtained from the Lunenfeld-Tanenbaum Research Institute’s OpenFreezer (Olhovsky et al. 2011). pcDNA5-miniTurbo-3xFLAG and pcDNA5-miniTurbo-3xFLAG-EGFP were gifts from Anne-Claude Gingras. pLVU/GFP and pLEX_305-N-dTAG lentiviral plasmid vectors were from Addgene (#24177, #91797). pHTN HaloTag CMV-neo and pHTC HaloTag CMV-neo vectors were from Promega (G7721, G7711). The coding sequences for FANCJ, RECQL5, GNB1L, TELO2, ATR were obtained from the ORFeome collection (http://horfdb. dfci.harvard.edu/), archived in OpenFreezer. The coding sequences for FANCJ, RECQL5, TELO2 were cloned into pHIV-NAT-T2A-hCD52 using NotI/XmaI restriction enzyme sites. The GNB1L coding sequence was cloned into pcDNA5-FRT/TO-3xFLAG, pcDNA5-miniTurbo-3xFLAG, pLVU/GFP, and pLEX_305-N-dTAG vectors using the Gateway system (Life Technologies/Thermo Fisher) according to the manufacturer’s protocol. The TELO2 and ATR coding sequences were cloned into pHTN HaloTag CMV-neo using SbfI/NotI restriction enzyme sites. The TELO2 coding sequence was cloned into pHTC HaloTag CMV-neo using SbfI/PvuI restriction enzyme sites

### Antibodies

The following primary antibodies were used in this study at the indicated dilutions for immunoblot analysis (IB), immunofluorescence (IF), or fluorescence-activated cell sorting (FACS): rabbit anti-H2A.X (1:1000 for IB; Cell Signaling, 2595), mouse anti-γ-H2AX JBW301 (1:1000 for IB, 1:2500 for IF, 1:5000 for FACS; Millipore, 05-636), mouse anti-γ-H2AX-Alexa Fluor 647 (1:1000 for FACS; Millipore, 05-636-AF647), rabbit anti-γ-H2AX (1:200 for IF; Cell Signaling, 2577), rabbit anti-53BP1 (1:2500 for IF, Santa Cruz, sc22760), mouse anti-RPA2(1:500 for IB and IF, Abcam, ab2175), rabbit anti-pRPA2 (S33) (1:1000 for IB; Bethyl, A300-246A-3), mouse anti-CHK1 (1:1000 for IB; Santa Cruz, sc8408), rabbit anti-pCHK1 (S345) (1:1000 for IB; Cell Signaling, 2348), mouse anti-PCNA (1:200 for IB; Santa Cruz, sc56), rabbit anti-ubiquityl PCNA (Lys164) (1:1000; Cell Signaling, 2577), rabbit anti-GNB1L (1:250 for IB; Millipore, HPA034627), rabbit anti-TELO2 (1:1500 for IB; Proteintech, 15975-1-AP), rabbit anti-TTI1 (1:3000 for IB; Bethyl, A303-451A), rabbit anti-TTI2 (1:1500 for IB; Bethyl, A303-476A), rabbit anti-RUVBL1 (1:2000 for IB; Proteintech, 10210-2-AP), rabbit anti-RUVBL2 (1:500 for IB; Abcam, ab36569), rabbit anti-PIH1D1 (1:1000 for IB; Proteintech, 19427-1-AP), rabbit anti-RPAP3 (1:1000 for IB; Bethyl, A304-854A-T), goat-anti-ATR (1:200 for IB; Santa Cruz, sc1887), goat-anti-DNA-PKcs (1:500 for IB; Santa Cruz, sc1552), rabbit anti-ATM (1:1000 for IB; Cell Signaling, 2873), rabbit anti-mTOR (1:2000 for IB; Cell Signaling, 2983), rabbit anti-TRRAP (1:2000 for IB; Bethyl, A301-132A), rabbit anti-SMG1(1:2000 for IB; Bethyl, A300-394A), mouse anti-α-Actinin (1:1000 for IB; Millipore, 05-384), mouse anti-α-tubulin (1:1000 for IB; Calbiochem, CP06), rat-anti-HA (1:1000 for IB; Roche, ROAHAHA), rabbit anti-RECQL5 (1:1000 for IB; Abcam, ab91422), rabbit anti-FANCJ (1:1000 for IB; MAB6496).

These secondary antibodies were used in 1:10000 dilution in immunoblot analysis: HRP-Sheep monoclonal anti-mouse IgG (Cytiva, #45-000-692); HRP-Goat polyclonal anti-rabbit IgG (Jackson Immunoresearch Labs, #111-035-144); HRP-Bovine polyclonal anti-goat IgG Jackson Immunoresearch Labs, #805-035-180). These secondary antibodies were used in 1:1000 dilution in immunofluorescence analysis: Alexa Fluor 488-Donkey polyclonal anti-rat IgG; Alexa Fluor 488-Donkey polyclonal anti-mouse IgG; Alexa Fluor 647-Donkey polyclonal anti-mouse IgG; Alexa Fluor 647-Donkey polyclonal anti-rabbit IgG (Thermo Fisher A-21208, A-21202, A-31571, A-31573).

### Immunofluorescence microscopy

To analyze γ-H2AX and 53BP1 focus formation in RKO *TP53^−/−^* cell lines, cells were seeded on coverslips to grow for 24 h, and then subjected to the indicated treatments or left untreated. Cells were rinsed with PBS once, subsequently fixed with 4% paraformaldehyde (PFA, Thermo Fisher) for 15 min at room temperature, and permeabilized with 0.3% Triton X-100 (Sigma, T8787) for 30 min. After fixation, cells were rinsed with PBS for three times, blocked in blocking buffer (10% goat serum (Sigma, G6767), 0.5% NP-40 (Sigma-Aldrich, I3021), 5% w/v saponin (Sigma-Aldrich, 84510), diluted in PBS) for 30 min, incubated with primary antibodies (mouse anti-γ-H2AX JBW301 1:2500 and rabbit anti-53BP1 1:2500) diluted in blocking buffer for 2 h at room temperature. Cells were then washed three times in PBS for 5 min and stained with secondary antibodies (Alexa Fluor 488-conjugated goat anti-mouse IgG and Alexa Fluor 647-conjugated goat anti-rabbit IgG, 1:1000 in blocking buffer) and 0.5–0.8 μg/mL DAPI (4,6-diamidino-2-phenylindole, Sigma-Aldrich, D9542) for 1 h at room temperature. Cells were washed as above, mounted in Pro-Long Gold mounting medium (Life Technologies), and imaged using a Zeiss LSM780 laser scanning microscope with a 60X objective. Image analysis was performed using Columbus (PerkinElmer) to quantify the nuclear foci of γ-H2AX and 53BP1 as described previously (Olivieri et al. 2020).

For immunofluorescence analysis of γ-H2AX and RPA2 in RKO *TP53^−/−^* cell lines, cells were grown on coverslips for 24 h, subjected to the indicated treatment, then pre-extracted for 10 min on ice with ice-cold buffer (25 mM HEPES, pH 7.4, 50 mM NaCl, 1 mM EDTA, 3 mM MgCl2, 300 mM sucrose and 0.5% Triton X-100) and fixed with 4% PFA for 15 min at room temperature. Staining was as described before except the primary antibodies used were rabbit anti-γ-H2AX 1:200 and mouse anti-RPA2 1:500. Images were acquired on a Zeiss LSM780 laser scanning microscope with a 20X objective and analyzed by Columbus (PerkinElmer) to quantify the nuclear intensity of γ-H2AX and RPA2 signals.

### Immunofluorescence and flow cytometry

Flow cytometry experiments were performed as described previously (Zimmermann et al. 2018). Briefly, cells were plated on 6-cm dishes to grow for 24 h before adding drugs. After drug treatment, cells were collected by trypsinization and centrifuged in a conical tube. Pellets were washed in PBS once and fixed in 4% PFA for 10 min at room temperature. Cells were spun, resuspended in 100 μl PBS and chilled on ice for 1 min. 900 μl of −20°C methanol was then added dropwise while gently vortexing. Fixed cells were stored at −20°C overnight or longer. Before staining, cells were spun down, washed with PBS, and blocked in blocking buffer (see “Immunofluorescence microscopy” section) at room temperature for 30 min. Cells were then centrifuged and resuspended in diluted Alexa Fluor 647-conjugated mouse anti-γ-H2AX antibody (Millipore, 05-636-AF647, 1:1000 in blocking buffer). After 2 h incubation the antibody was diluted with 10X volume PBS, cells were spun down and resuspended in PBS with DAPI. Cells were analyzed on BD LSRFortessa X-20 (BD Biosciences), or MoFlo Astrios EQ Cell Sorter (Beckman Coulter), or Attune NxT/CytKick Max autosampler (Thermo Fisher).

### Phenotypic CRISPR/Cas9 screens based on γ-H2AX

RKO *TP53^−/−^* cells or COL-hTERT *TP53^−/−^* cells were transduced with the lentiviral TKOv3 library (Hart et al., 2017) at a low MOI (∼0.3) and puromycin-containing medium was added the next day. Three days after transduction, which was considered the initial time point (T0), cells were pooled together and divided in two technical replicates (the only exception is the untreated RKO screen in which we did four replicates). Each replicate was cultured for five more days to provide time for sgRNA-mediated gene editing, then divided into different treatments at T5. Cells were either treated with 0.3 μM Aph, or with 0.2 μM Ara-C, or with 200 μM HU, or left untreated (UT) for 24 h. At T6, 40 million cells per sample were collected in pellets and frozen at −80 °C as the unsorted population, with the remaining cells (∼ 400 million) subjected to fixation, staining and FACS. These cells were spun down in 50 ml conical tubes, washed with PBS once, and fixed in 4% PFA for 10 min at room temperature while rotating. Cells were then pelleted and resuspended in 1 ml PBS and chilled on ice. 19 ml of −20°C methanol was then added dropwise while gently vortexing. Fixed cells were stored at −20°C overnight or longer. Before staining, cells were spun down, washed with FACS buffer (PBS with 5% FBS), and blocked in blocking buffer at room temperature for 30 min while rotating. Cells were then centrifuged and resuspended in diluted Alexa Fluor 647-conjugated mouse anti-γ-H2AX antibody (Millipore, 05-636-AF647, 1:1000 in blocking buffer). After 2 h incubation,40 ml of FACS buffer was added, cells were spun down and resuspended in 10 ml PBS with DAPI, then subjected to sorting on a MoFlo Astrios EQ Cell Sorter. Cells with the top 5% of γ-H2AX signal intensity were collected and their genomic DNA (gDNA) was extracted using the FFPE DNA Purification Kit (Norgen Biotek Cat. 47400). gDNA from unsorted cell population was isolated using the QIAamp Blood Maxi Kit (Qiagen). For both sorted and the unsorted cell populations, genome-integrated sgRNA sequences were amplified by PCR using KAPA HiFi HotStart ReadyMix (Kapa Biosystems). i5 and i7 multiplexing barcodes were added in a second round of PCR and final gel-purified products were sequenced on Illumina NextSeq500 systems to determine sgRNA representation in each sample.

### Drugs

The following drugs were used in this study: Aph (Focus Biochemicals, 10-2058), HU (Sigma, H8627), Ara-C (Sigma, C1768), CD437 (Sigma, C5865), gemcitabine (Cayman chemical, 9003096), dTAG^V^-1 (gift from Benham Nabet), TBB (4,5,6,7-tetrabromobenzotriazole, Selleckchem, S5265).

### Immunoprecipitation

Cells were transfected with pcDNA5-3xFLAG-GNB1L or pcDNA5-3xFLAG-TELO2. 48 hours later, cells were collected by trypsinization, washed with PBS once, and lysed in 1 ml high salt lysis buffer (50 mM HEPES pH8, 300 mM NaCl, 2 mM EDTA, 0.1% NP-40, 10% glycerol, plus protease inhibitors (cOmplete EDTA-free protease inhibitor cocktail, Roche, 11836170001)). Lysates were incubated with gentle rotation at 4°C for 30 min with occasional vortexing and then centrifuged at 15,000xg for 10 min. 150 μl of total cell lysates were used as input and 850 µl were incubated with 40 μl anti-FLAG M2 magnetic bead (Sigma M8233) at 4°C overnight while rotating. Beads were washed three times with TBS (50 mM Tris-HCl pH 7.4, 150 mM NaCl) and eluted with 100 μl of 3x FLAG peptide (100 μg/ml, GLPBio, GP10149) at 4°C for 30 min. Elution was repeated once more and 40 μl 6x SDS-PAGE sample buffer was added to each sample. Samples were boiled at 95°C for 5 min and subjected to SDS-PAGE and immunoblot analysis.

### Parallel miniTurbo proximity labeling and affinity purification (AP) coupled to mass spectrometry (MS)

Parental 293 Flp-In cells, and cells stably expressing miniTurbo-3xFlag-GNB1L, miniTurbo-3xFlag-eGFP or miniTurbo-3xFlag were used for parallel miniTurbo and AP-MS studies. For both miniTurbo and AP-MS, two 150-mm plates of cells were treated with 5 μg/ml doxycycline for 24 h to induce expression of bait proteins. For miniTurbo, 50 μM biotin was added to cells 40 min before harvest. Cells were pelleted at low speed, washed with ice-cold PBS and frozen at −80°C until purification. Cell lysis, purification, and mass spectrometry were performed as previously described in (Samavarchi-Tehrani et al. 2018).

### Incucyte cell growth assay

For Aph dose-response assays in RKO *TP53^−/−^* parental (WT) and *FANCJ-KO* cells, 770 cells per well were seeded in 96-well plates and treated with sequential serial dilutions of Aph. After 6 days of treatment, the cell confluency was measured using an IncuCyte Live-Cell Analysis system (Sartorius). Confluence growth inhibition was calculated as the relative confluency compared to untreated cells. For proliferation assays of sg*DERA-* and sg*GNB1L*-expressing RKO *TP53^−/−^* cells, 6000 cells per well were seeded in 24-well plates and treated with the indicated replication inhibitor or left untreated. The cell confluency was measured once 24 h post-seeding using an IncuCyte Live-Cell Analysis system. Growth curves were generated using confluency as the proxy for cell numbers.

### dTAG-mediated protein degradation system

Two clonal 293T cell lines were generated by lentiviral transduction to introduce sg*GNB1L*-1 and the sgRNA-resistant *FKBP_mut_-GNB1L* plasmid into the parental 293T cells. The dTAG^V^-1 compound was added at 1 μM for 1-24 h (short-term) or 1-6 days (long-term) to induce the degradation of the FKBP_mut_-GNB1L protein (Nabet et al. 2018; Nabet et al. 2020).

### HaloTag label-switch experiments

293T cells were transiently transfected with the Halo-ATR plasmid 24 h before the labeling experiments. To label newly synthesized Halo-ATR, cells were incubated with 10 μM of the blocking agent 7-bromoheptanol (Merrill et al. 2019) (Alfa Aesar, H54762), for 2 h, followed by two washes, and then incubated with 1 μM TMR HaloTag ligand (Promega, G8252) for indicated time. To label pre-existing Halo-ATR, cells were incubated with 1 μM TMR HaloTag ligand for 1 h, followed by two washes, and then incubated with 10 μM blocking agent for the indicated time. Whole cell lysates were analyzed by SDS-PAGE and immunoblotting. TMR fluorescence signal was measured with a Typhoon FLA 9500 laser scanner (GE Healthcare). ImageJ (imagej.nih.gov) was used to quantify band intensities of TMR and α-actinin.

### POPcode mutagenesis screen

The *GNB1L* open reading frame (ORF) was inserted into the pLVU/GFP lentiviral plasmid vector (Addgene, 24177) encoding a C-terminal GFP tag. The *GNB1L* coding sequence was subdivided into two regions, and variant libraries were generated via the oligonucleotide-directed mutagenesis method POPCode (Weile et al. 2017; Sun et al. 2020). Both *GNB1L* variant libraries were introduced into RKO *TP53^−/−^* cells separately by lentiviral transduction. Cells were sorted for the GFP-positive population to select for *GNB1L-GFP* variant integration. Cells were then transduced with sg*GNB1L*-1 (Supplementary Table 8) which targeted at an intron-exon junction site of the *GNB1L* gene. The γ-H2AX FACS screen was performed in cells with *GNB1L-GFP* variant library/sg*GNB1L*-1 in the presence of 150 μM HU, as described previously in the “Phenotypic CRISPR/Cas9 screen” section. Cells with the top 5% of γ-H2AX signal intensity were collected, and genomic DNA was extracted from both sorted and unsorted cells. The primer set (forward: 5’ TCTGGCCGTTTTTGGCTTTTT 3’; reverse: 5’ GAACAGCTCCTCGCCCTTG 3’) was used for PCR amplification of the inserted *GNB1L* ORF sequence. Variant frequencies in the pre- and post-selection libraries were assessed using TileSeq (Weile et al. 2017; Sun et al. 2020). Briefly, each ‘tile’ within the target locus was amplified with primers including Illumina sequencing adapters, followed by the addition of Illumina indices in a low-cycle PCR. Tiled libraries (including a wild type control) were sequenced by paired-end sequencing on an Illumina NextSeq 500 device using 300 cycle NextSeq 500/550 Mid Output v2.5 Kits, generating ∼2M reads per tile. Sequencing data was processed as described previously (Weile et al. 2017; Sun et al. 2020). Briefly, libraries were demultiplexed with Illumina bcl2fastq and variant allele frequencies for each condition were calculated using the tileseq-package. Here, reads are aligned to a template sequence and mutations are called where there is agreement between both forward and reverse reads. Where read pairs disagreed, variants were treated as wild type. Fitness values were calculated using the tileseqMave pipeline (Weile et al. 2021) and scores were scaled based on the distribution of synonymous variants and the bottom 5^th^ percentile of functional scores (in the absence of nonsense variants).

### AlphaFold2 prediction of GNB1L-TELO2 interaction

Amino acid sequences corresponding to full-length human GNB1L and TELO2(460-640) were used as two separate chain inputs for the ColabFold implementation of AlphaFold2-multimer (Evans et al. 2022; Mirdita et al. 2022) using the following parameters: no templates, no amber relax, MMSeqs2 MSA mode, AlphaFold2-multimer-v2 model type, 5 models, 3 recycles. The top-ranking model was used for further analysis. The predicted aligned error plot for the top ranking model is shown in Fig S10C. A plot displaying the mean interface predicted aligned error (PAE) is shown in Fig S10D. Mean interface PAE is defined as the average PAE value between the indicated TELO2 residue and every GNB1L residue predicted to be within 9 Å. TELO2 residues without any nearby GNB1L residues are assigned the maximum PAE value. Molecule display and analysis were performed in ChimeraX (Pettersen et al. 2021).

### GO-Figure!

GO-Figure! analyses in Figures 1C and S1B were done using the python script provided by Waterhouse Lab, using the top 80 GO terms sorted by adjusted p-value, as calculated by Enricher (https://maayanlab.cloud/Enrichr/). The GO-figure software clusters GO terms together at a similarity threshold of 0.5, using a weighted distance algorithm to describe proximity in the GO hierarchy. The figure represents similarity of GO term clusters based on their hierarchical proximity, with point color describing adjusted p-value of the representative term, and point size describing the number of GO terms within each cluster.

### FoldX and rolling window analysis

ΔΔG scores were calculated using the BuildModel command of the FoldX software, following the methodology presented by ELELAB’s mutateX algorithm (Figure 6B, and Supplementary Tables 5 and 6). Each residue of the protein of interest was mutated to every other possible amino acid using the standard parameters of BuildModel.

The rolling window analysis plots (Figure 6B) take the rolling mean of each of three score types with a window size of five, therefore each position represents the mean of the given position along with the two scores before and after. The functional and ΔΔG scores were initially summarized for each position by taking the mean of all mutation scores for that residue, not including mutation to self or termination where it was provided. The conservation scores were calculated directly for each position with ConSurf (consurf.tau.ac.il/consurf_index.php).

### Quantification and statistical analysis

All data presented are biological replicates unless otherwise stated. The statistical tests used, number of replicates, definition of error bars and center definitions are all defined within each figure or figure legend. In Figures 2C, 2E, 3B, 3E, 4C, 5E, 6D, S4B, S4E, S4F, S5C, S5D, S6C-E, S7A, S8B, S8D, S9C, S9D, S10B, Student’s unpaired t-test was used to calculated mean, standard deviations, and p-values in GraphPad Prism 9. ns = p > 0.05, * = p < 0.05, ** = p < 0.01, *** = p < 0.001. In Figure 3D, nonlinear fitting of drug dose response curves was also performed using GraphPad Prism 9.

## Acknowledgements

We thank Rachel Szilard for critical reading of the manuscript and Junjie Chen for sharing results prior to their publication. We also thank Laurence Pearl, Mohinder Pal and Oscar Llorca for discussions on the TTT complex, Agnel Sfeir for RKO *TP53^−/−^* cells and Benham Nabet for sharing dTAG^V^-1. YZ is supported by an Ontario Trillium Scholarship. Work in the DD lab was supported by grants from the Canadian Cancer Society (705644) and Canadian Institutes for Health Research (CIHR, grant FDN143343) with additional support from the Krembil Foundation.

## Author Contributions

Yichao Zhao: Conceptualization, Investigation, Writing and Visualization; Daniel Tabet: Investigation, Formal Analysis; Diana Rubio Contreras: Investigation; Arne Nedergaard Kousholt: Resources; Jochen Weile: Formal analysis; Henrique Melo: Formal Analysis; Lisa Hoeg: Formal Analysis, Data Curation, Visualization; Atina G. Coté: Investigation; Zhen-Yuan Lin: Investigation; Dheva Setiaputra: Investigation, Visualization; Jos Jonkers, Supervision, Resources; Fernando Gómez Herreros: Supervision; Frederick P. Roth: Conceptualization, Supervision, Resources; Daniel Durocher: Conceptualization; Supervision, Writing, Funding acquisition.

## Conflict of interest statement

DD is a shareholder and advisor for Repare Therapeutics

---

**Figure S1.**
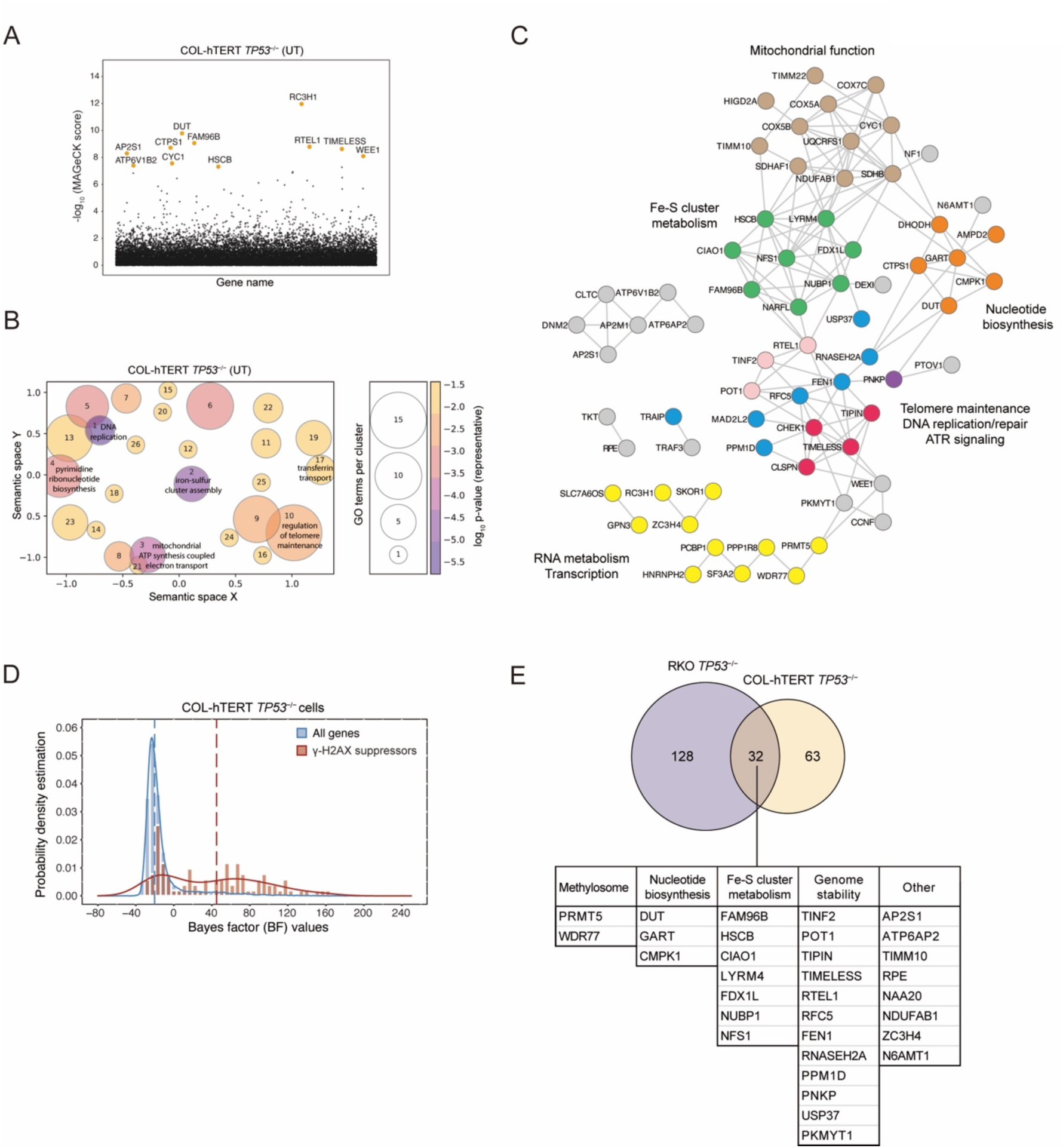
Genes and pathways that prevent endogenous DNA damage in COL-hTERT *TP53^−/−^* cells. (A) Manhattan dot plots of γ-H2AX screen results in untreated (UT) COL-hTERT *TP53^−/−^* cells. The top 11 genes were highlighted. (B) Gene Ontology (GO) analysis of Biological Process was performed using Enrichr for 95 γ-H2AX suppressors in COL-hTERT *TP53^−/−^* cells. The top 80 GO terms ranked by p-value were visualized by GO-Figure! software using a similarity cutoff of 0.5. Color represents the p-value of the representative GO term for the cluster, and the size of the circle represents the number of GO terms in a cluster. (C) STRING network analysis of γ-H2AX suppressors in COL-hTERT *TP53^−/−^* cells, based on information from text mining, experiments, databases, and neighborhood, with a high confidence threshold. Out of 95 genes, 60 are mapped to the network. Pathways of genes are manually curated with different colors: green, Fe-S cluster assembly; orange, nucleotide biosynthesis; brown, mitochondrial function; yellow, RNA metabolism; pink, telomere maintenance; purple, DNA repair; blue, DNA replication; red, ATR signaling; grey, others. (D) Distributions of gene essentiality scores of γ-H2AX suppressors (brown) and whole genome reference (blue) in COL-hTERT *TP53^−/−^* cells. Dashed lines indicate the median for each population. (E) Venn diagram of γ-H2AX suppressors in RKO *TP53^−/−^* and COL-hTERT *TP53^−/−^* cells with the list of γ-H2AX suppressors common to the RKO *TP53^−/−^* and COL-hTERT *TP53^−/−^* cell lines. Pathways of genes were annotated manually.

**Figure S2.**
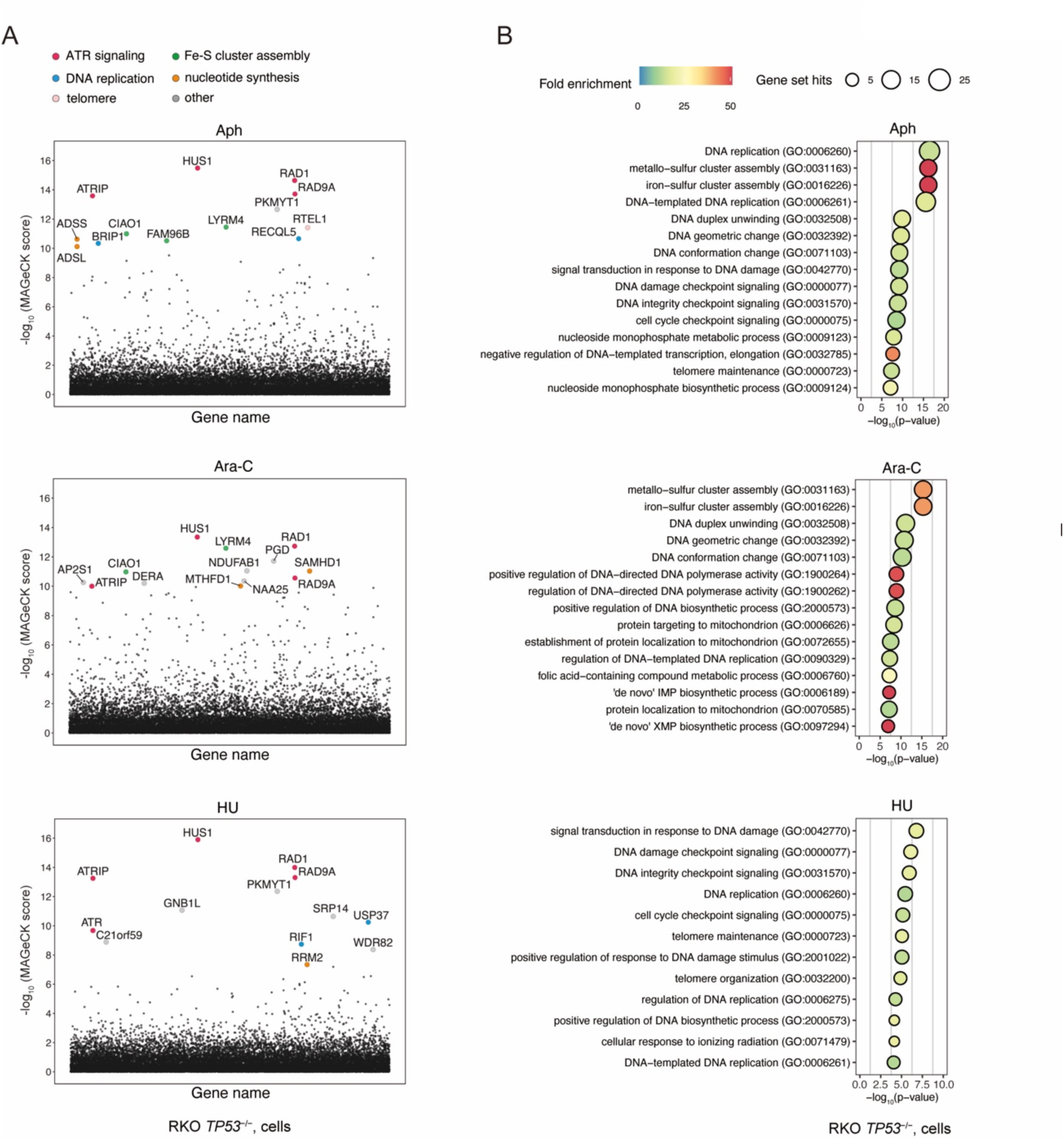
Genes and pathways that prevent replication-associated DNA damage in RKO *TP53^−/−^* cells. (A) Manhattan dot plots of γ-H2AX screen results in RKO *TP53^−/−^* cells treated with Aph (Aph), Ara-C (Ara-C), HU (HU). Top 13 genes from each screen were highlighted. (B) Gene Ontology term enrichment analysis of Biological Process of the gene hits (FDR<0.05) in RKO *TP53^−/−^* Aph, Ara-C, and HU screens, using the Fisher exact test with FDR correction. Circle size indicates the number of genes included in each GO term, color indicates the fold enrichment compared to the whole genome reference set, and x-axis position indicates negative log_10_ FDR value.

**Figure S3.**
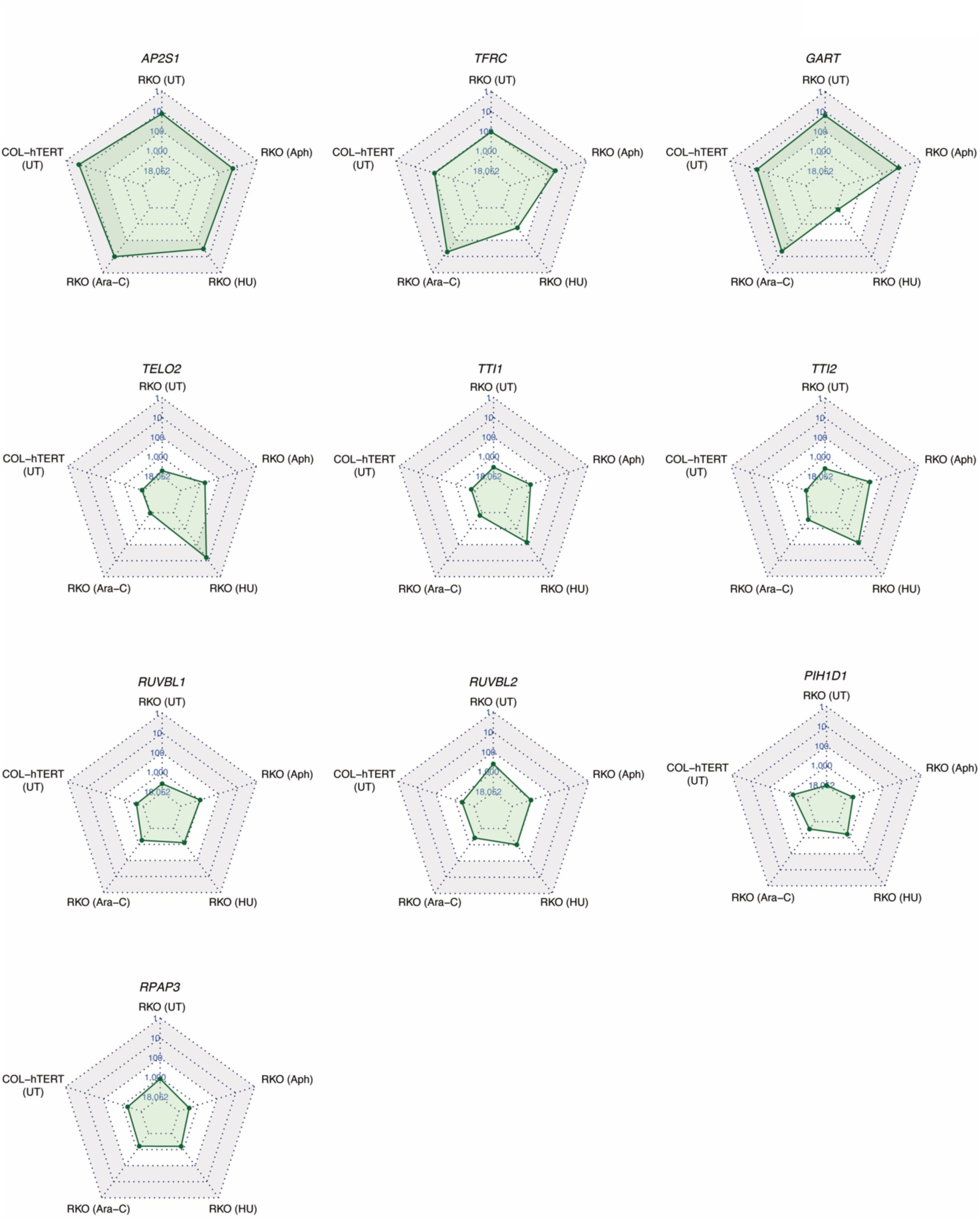
Radar plots of gene ranking in five γ-H2AX screens. Radar plots showing the ranking of *AP2S1, TFRC, GART, TELO2, TTI1, TTI2, RUVBL1, RUVBL2, PIH1D1,* and *RPAP3* in all five γ-H2AX screens.

**Figure S4.**
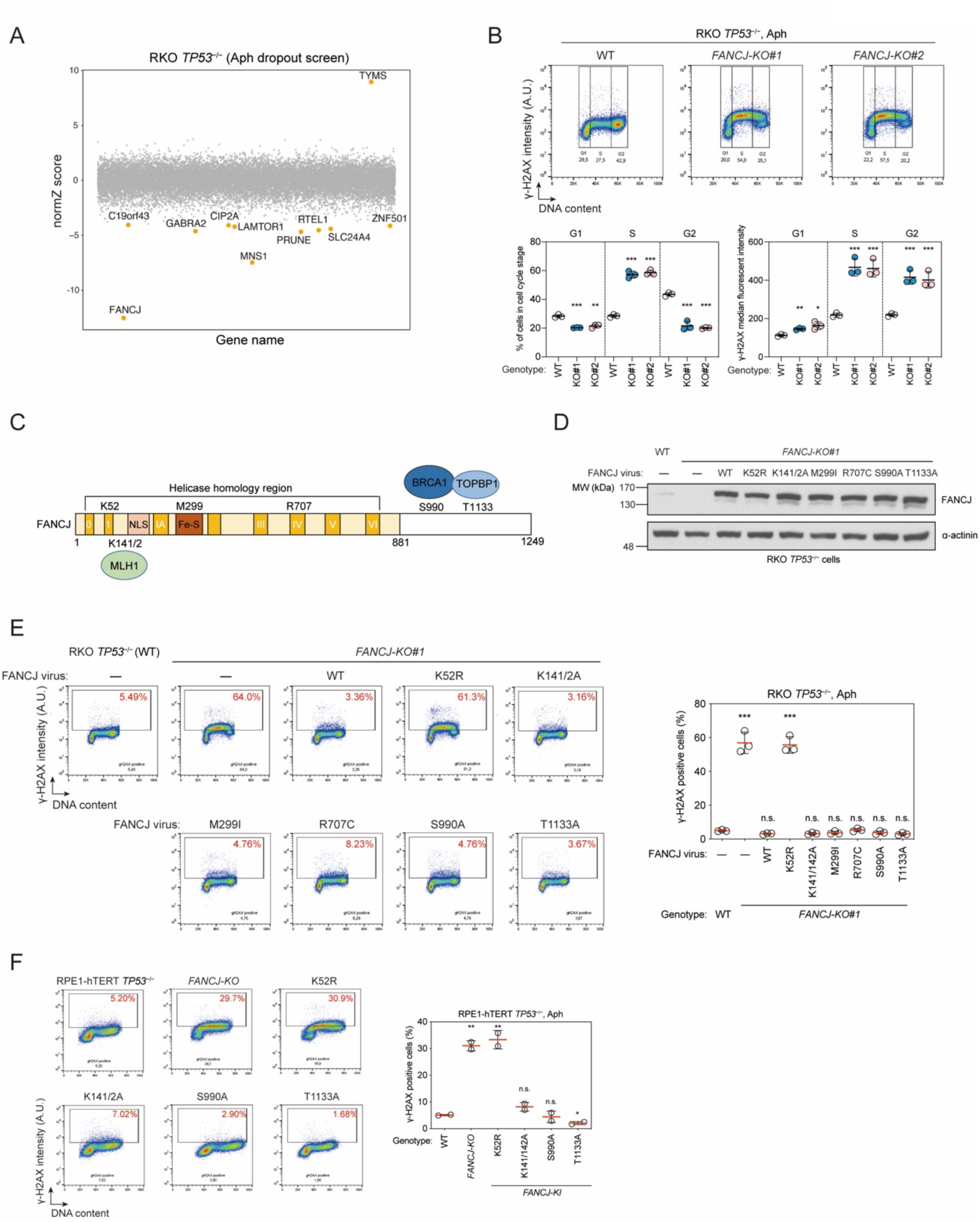
Characterization of *FANCJ-KO* cells. Related to Figure 3. (A) Manhattan dot plots of the CRISPR/Cas9 fitness screen results in RKO *TP53^−/−^* cells with Aph treatment. The most Aph-resistant gene *TYMS* and top 11 Aph-sensitizing genes are highlighted. (B) Flow cytometry analysis of RKO *TP53^−/−^* parental (WT) and *FANCJ-KO* cells. Cells were treated with 300 nM Aph for 24 hours, then fixed and subjected to staining with a γ-H2AX antibody and DAPI. Upper panel, representative flow cytometry plots. Lower panel, quantification of number of cells (left) and γ-H2AX intensity (right) in G1, S, and G2 phases. Cell cycle gating was based on DNA content (DAPI). Bars represent the mean ± s.d. (n=3 independent experiments). Results of an unpaired t-test between WT and *FANCJ-KO* cells are shown (*p<0.05; **p<0.01; *** p<0.001). (C) Schematic representation of the FANCJ protein. Helicase domains 0, I, IA, II, III, IV, V, VI; NLS, nuclear localization sequence; Fe-S, iron-sulfur cluster binding domain. MLH1, BRCA1 and TOPBP1 are known binding partners of FANCJ. (D) Immunoblot analysis of FANCJ expression in RKO *FANCJ-KO#1* cells complemented with wildtype (WT) or mutant *FANCJ* variants as indicated using lentiviral transduction. (E) Flow cytometry analysis of the cells described in (D), treated with 300 nM Aph. Left, representative flow cytometry plots. Right, quantification of γ-H2AX positive cells. Bars represent the mean ± s.d. (n=3 independent experiments). Comparisons are made to WT cells transduced with an empty vector, using an unpaired t-test (***p<0.001; n.s., not significant). (F) Flow cytometry analysis of RPE1-hTERT *TP53^−/−^* cells of the indicated genotype. Bars represent the mean ± s.d. (n=3 independent experiments). Comparisons are made to the WT *FANCJ* cells, using an unpaired t-test (*p<0.05; **p<0.01; n.s., not significant). A.U., arbitrary units.

**Figure S5.**
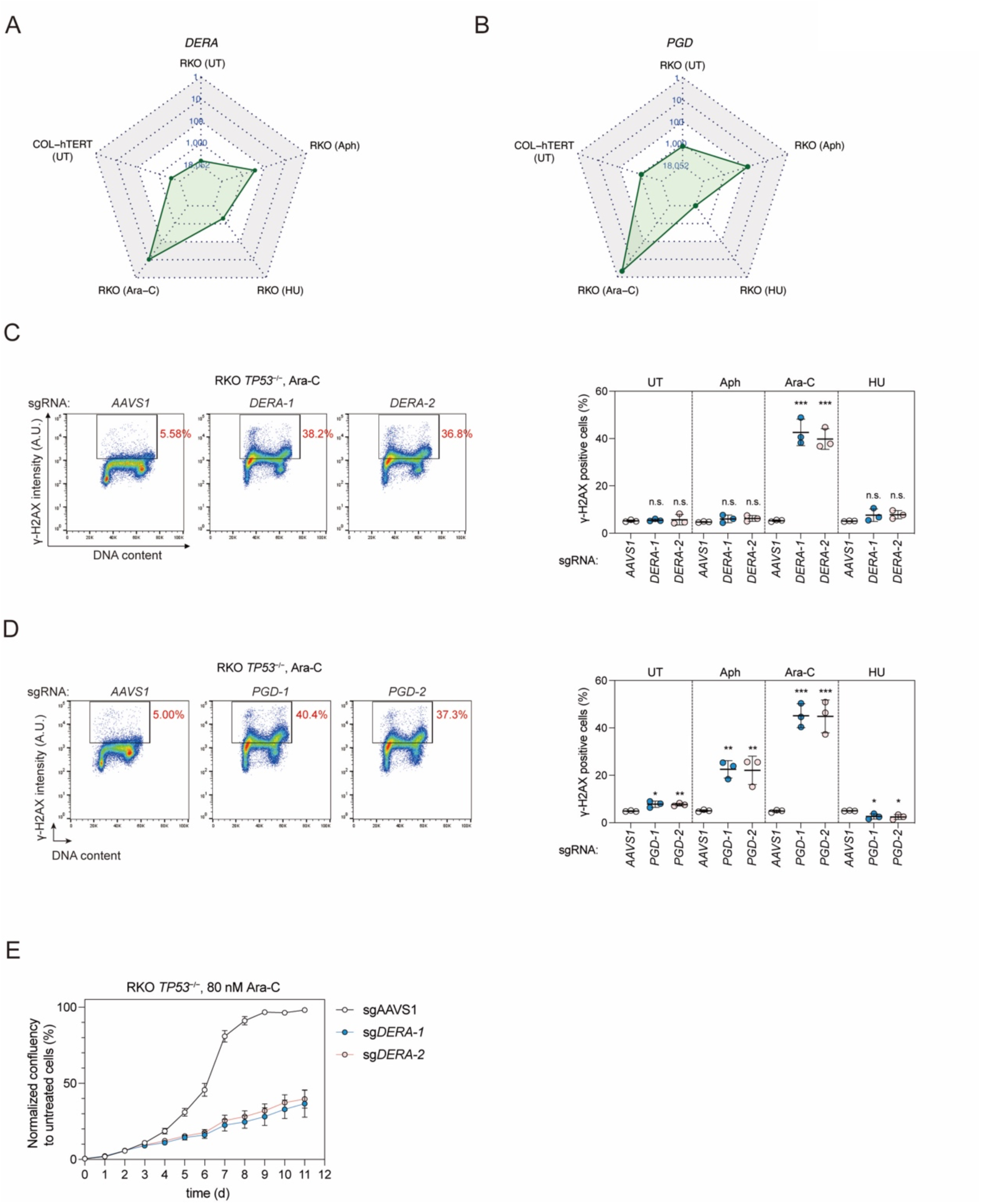
*DERA* and *PGD* act against Ara-C-induced replication stress. (A), (B) Radar plots showing the ranking of (A) *DERA* and (B) *PGD* in all five γ-H2AX screens. (C), (D) Flow cytometry analysis of RKO *TP53^−/−^* cells expressing the indicated sgRNA. Cells were treated with the indicated replication inhibitor for 24 h or left untreated. Cells were then fixed and stained with a γ-H2AX antibody and DAPI. Left, representative flow cytometry plots of cells treated with Ara-C. Right, quantification of γ-H2AX positive cells in all four conditions. Bars represent the mean ± s.d. (n=3 independent experiments). Comparisons are made to the sg*AAVS1* control within each treatment condition, using an unpaired t-test (*p<0.05; **p<0.01; ***p<0.001; n.s., not significant). A.U. arbitrary units. (E) Proliferation curves of RKO *TP53^−/−^* cells expressing sg*AAVS1* control or sg*DERA* in the presence of 80 nM Ara-C. Confluency is normalized to the untreated condition for each genotype. Data is presented as mean ± s.d. (n=3 independent experiments).

**Figure S6.**
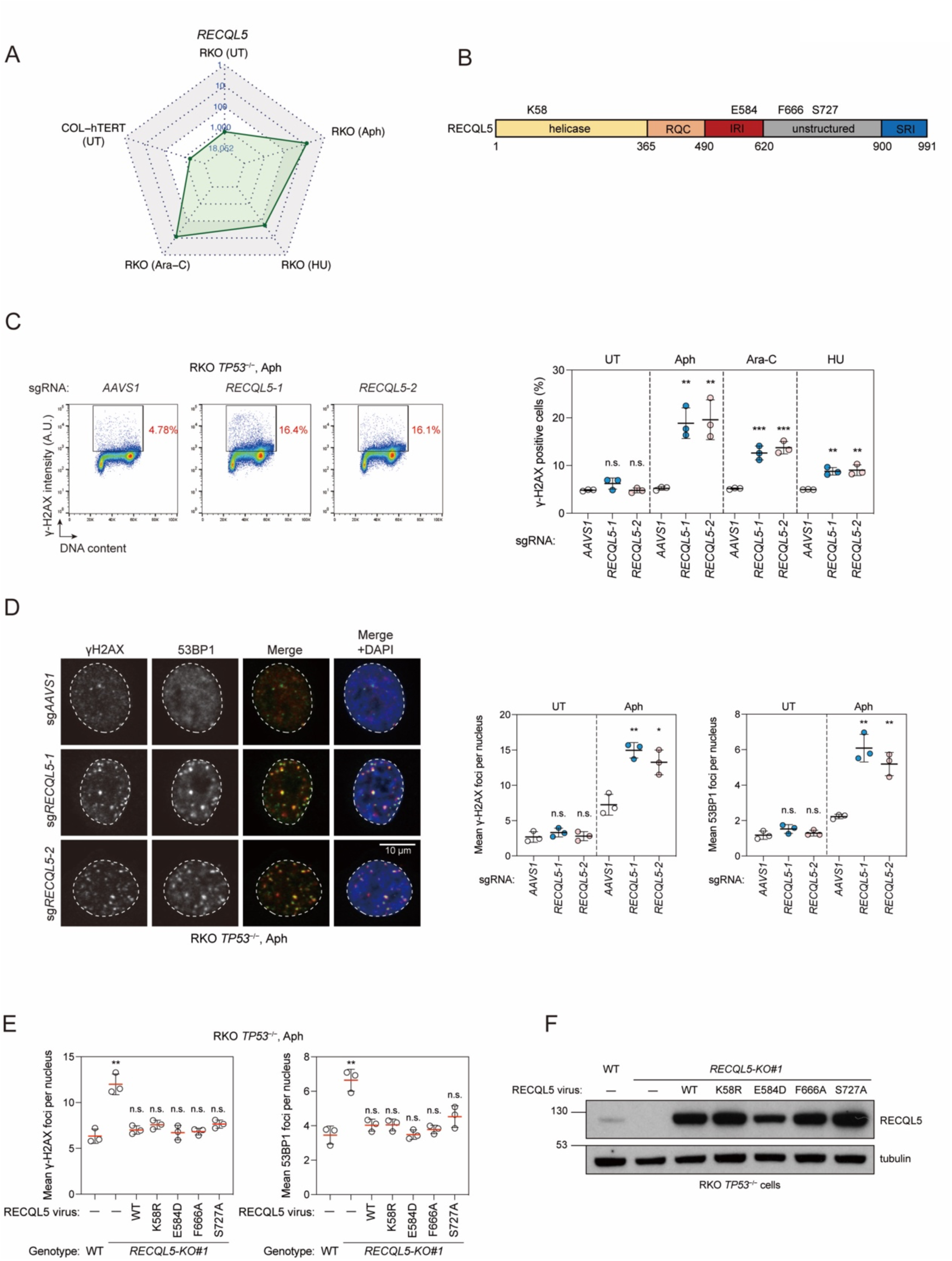
*RECQL5* protects cells from replication-associated DSBs. (A) A radar plot showing the ranking of *RECQL5* in all five γ-H2AX screens. (B) Schematic representation of the RECQL5 protein. RQC, RecQ carboxy–terminal domain; IRI, internal Pol II interaction domain; SRI, Set2–Rpb1– interacting domain. (C) Flow cytometry analysis of sgRNA-transduced RKO *TP53^−/−^* cells grown in the presence or absence of replication inhibitors. Left, representative flow cytometry plots of cells treated with Aph. Right, quantification of γ-H2AX positive cells in all four conditions. Bars represent the mean ± s.d. (n=3 independent experiments). Comparisons are made to the sg*AAVS1* control within each treatment condition, using an unpaired t-test (**p<0.01; ***p<0.001; n.s., not significant). (D) Immunofluorescence analysis of sgRNA-transduced RKO *TP53^−/−^* cells using γ-H2AX and 53BP1 antibodies. Left, representative images of three immunostainings. Dashed lines indicate the nuclear area determined by DAPI staining. Right, quantification of mean γ-H2AX and 53BP1 focus numbers. Each experiment includes a minimum of 500 cells for analysis. Bars represent the mean ± s.d. (n=3 independent experiments). Comparisons are made to the sg*AAVS1* control, using an unpaired t-test (*p<0.05; **p<0.01, n.s., not significant). (E) Immunofluorescence analysis of RKO *TP53^−/−^* cells of the indicated genotype transduced with an empty lentivirus (−) or one encoding WT RECQL5 or the indicated variants. Cells were treated with 300 nM Aph for 24 h before fixation and staining with antibodies to γ-H2AX and 53BP1. Mean γ-H2AX and 53BP1 focus numbers were quantified. Bars represent the mean ± s.d. (n=3 independent experiments). Comparisons are made to WT cells with empty vector, using an unpaired t-test (*p<0.05; **p<0.01, n.s., not significant). (F) Immunoblot analysis of RECQL5 expression in the cells described in E. Tubulin, loading control.

**Figure S7.**
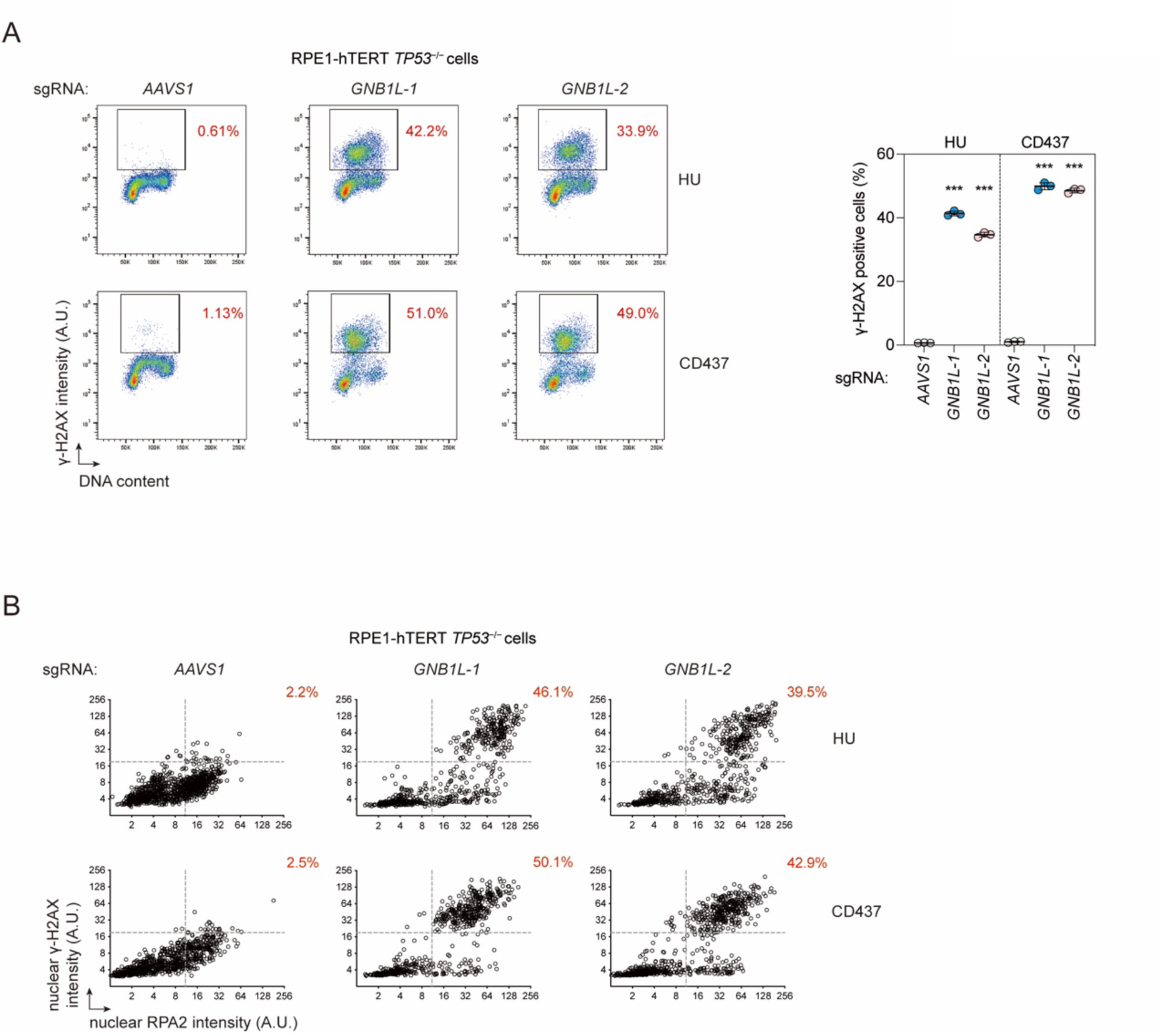
*GNB1L* protects RPE1-hTERT *TP53^−/−^* cells from replication catastrophe under mild replication stress. Related to Figure 4. (A) Left, representative flow cytometry analysis of RPE1-hTERT *TP53^−/−^* cells expressing sg*AAVS1* control or sg*GNB1L*. Cells were treated with 200 μM HU or 250 nM CD437 for 24 hours, then fixed and stained with a γ-H2AX antibody and DAPI. Red numbers indicate the percentage of γ-H2AX positive cells. Right, quantification of the experiment shown on left. Bars represent the mean ± s.d. (n=3 independent experiments independent experiments). Comparisons are made to the sg*AAVS1* control within each treatment condition, using an unpaired t-test (*** p<0.001). (B) Quantitative image-based cytometry (QIBC) analysis of γ-H2AX and chromatin-bound RPA2 signal intensities in RPE-hTERT *TP53^−/−^*cells, as described in (A). Cells were treated with 200 μM HU or 250 nM CD437 for 24 hours, then extracted, fixed and stained with antibodies to γ-H2AX and RPA2. Red numbers indicate the percentage of cells with both high γ-H2AX and high RPA2 signal intensities. A.U., arbitrary units.

**Figure S8.**
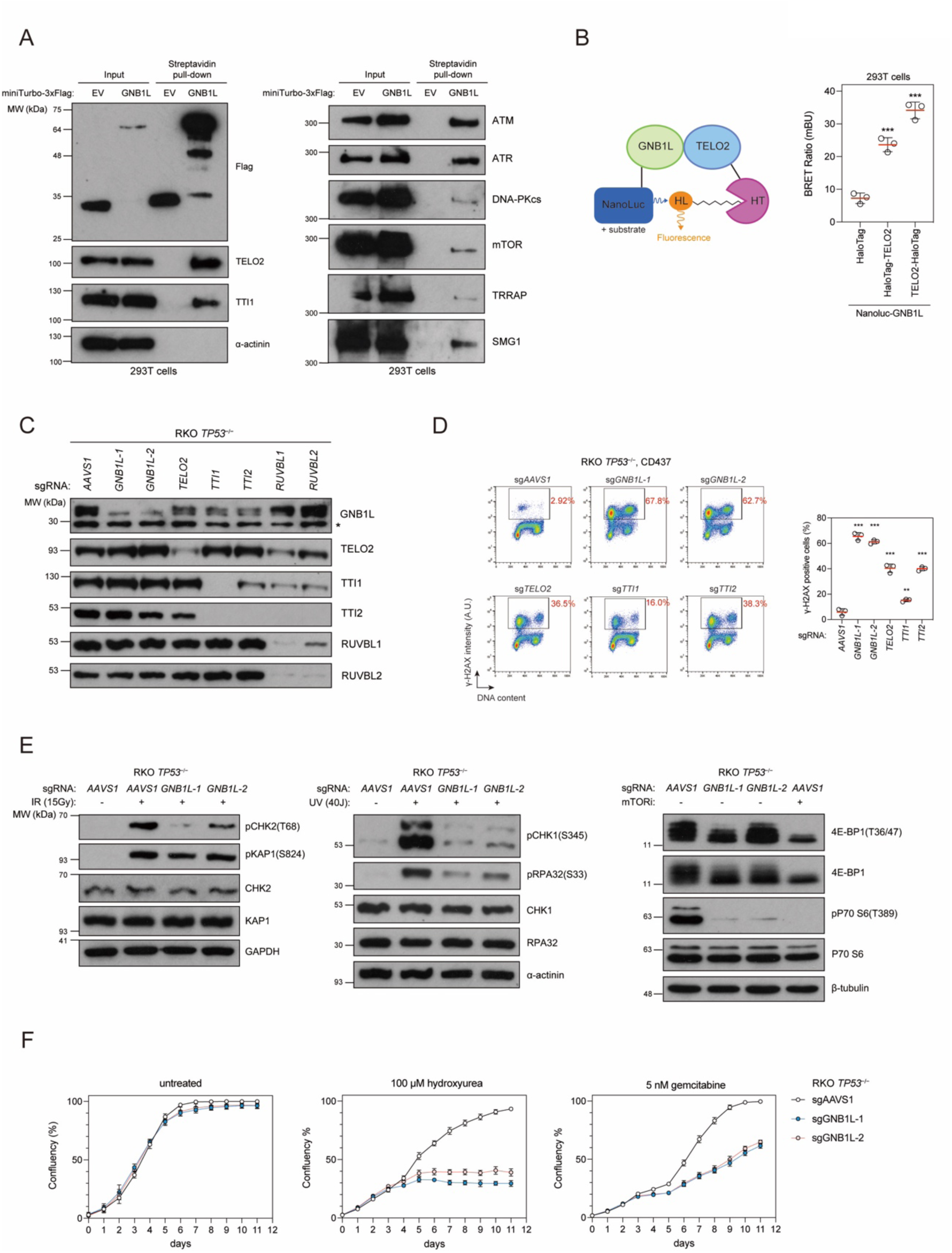
GNB1L associates with PIKKs and is essential for the downstream signaling of ATM/ATR/mTOR. Related to Figure 5. (A) Streptavidin pull-down was performed using lysates of 293T cells expressing miniTurbo-GNB1L or miniTurbo alone. 50 μM biotin was added to the cell culture medium 40 min before harvesting. Immunoblot analysis was performed with the indicated antibodies. (B) NanoBRET assay to assess the in vivo interaction between GNB1L and TELO2. Bars represent the mean ± s.d. (n=3 independent experiments). Results of unpaired t-test between unfused HaloTag and HaloTag TELO2 conditions are shown (*** p<0.001). (C) RKO *TP53^−/−^* cells were transduced with lentiviruses expressing the indicated sgRNA and cell lysates were subjected to immunoblot analysis. α-actinin, loading control. Asterisk besides GNB1L immunoblots indicate non-specific bands. (D) Flow cytometry analysis of RKO *TP53^−/−^* cells expressing the indicated sgRNA. Cells were treated with 250 nM CD437 for 24 hours, and then fixed, stained with a γ-H2AX antibody and DAPI. Left, representative plots. Right, quantification of γ-H2AX positive cells. Bars represent the mean ± s.d. (n=3 independent experiments). Comparisons are made to the sg*AAVS1* control, using an unpaired t-test (***p<0.001; **p<0.01). A.U., arbitrary units. (E) Substrate phosphorylation of ATM, ATR, mTOR in RKO *TP53^−/−^* cells expressing sg*AAVS1* control and sg*GNB1L*. To test ATM activity, cells were exposed to 15 Gy ionizing radiation (IR) and allowed to recover for 1 h before harvesting. To test ATR activity, cells were exposed to 40 J of ultraviolet (UV) and allowed to recover for 5 h. mTOR inhibitor Torin 1 was added to cells at a final concentration of 200 nM for 24 hours as a control for decreased mTOR activity. (F) Proliferation curves of the cells described in (E) grown in the presence of 100 μM HU, or 5 nM gemcitabine, or in the untreated condition. Data is presented as mean ± s.d. (n=3 independent experiments).

**Figure S9.**
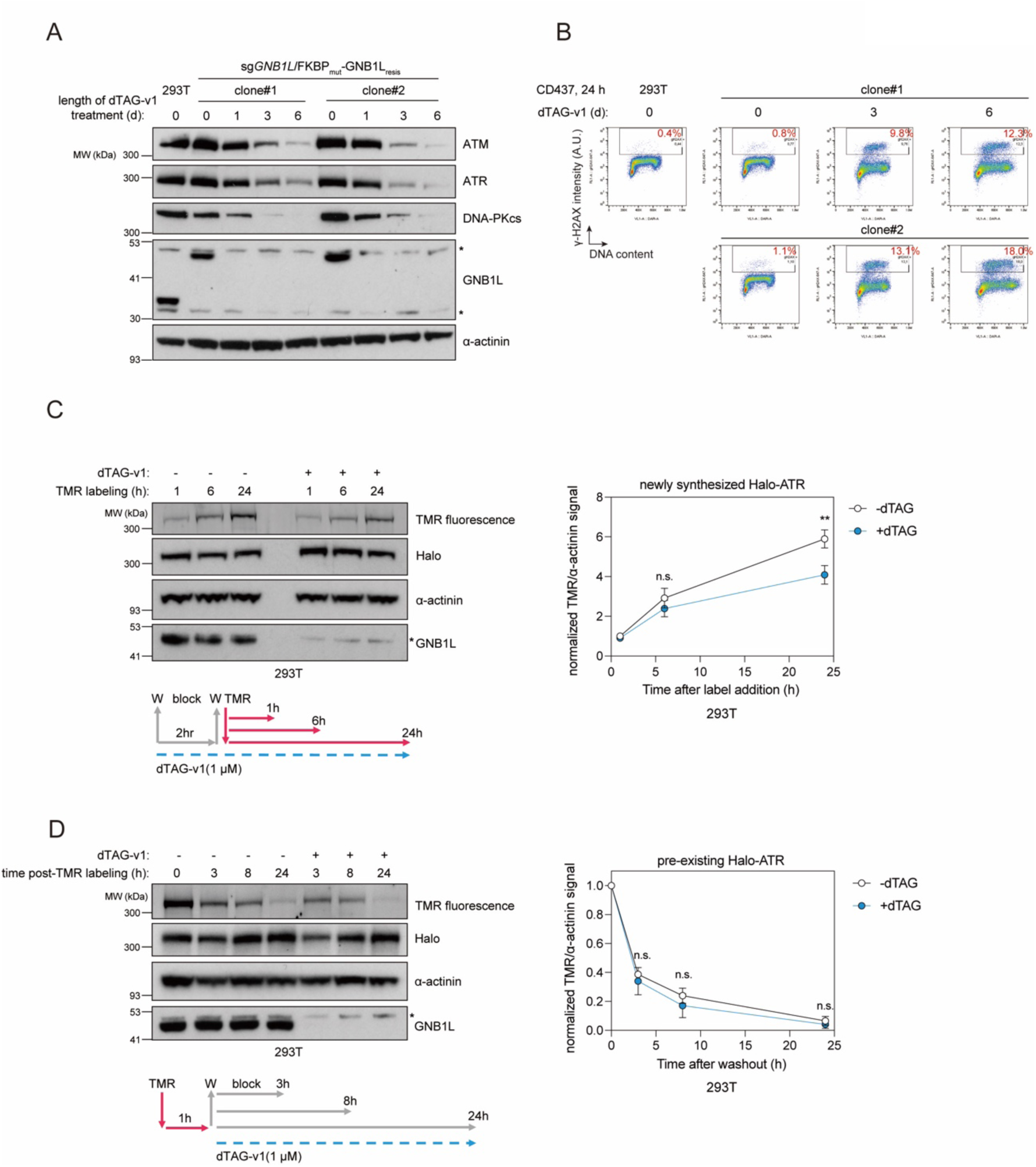
GNB1L maintains the protein level of newly synthesized but not pre-exsiting Halo-ATR. (A), (B) Lentiviral transduction of 293T cells were used to introduce a construct encoding sgRNA-resistant FKBP_mut_-GNB1L_resis_; degradation of this protein can be induced by addition of the dTAG-v1 molecule. Endogenous *GNB1L* was then inactivated with an sgRNA targeting an intron-exon junction. Two clones of the above-mentioned genotype were generated and were treated with 1 μM dTAG-v1 molecule for the indicated time before harvesting. (A) Immunoblot analysis of cell lysates with the indicated antibodies. (B) Flow cytometry analysis of cells treated with 250 nM CD437 for 24 hours. Cells were stained with a γ-H2AX antibody and DAPI. Red numbers indicate the percentage of γ-H2AX positive cells. (C), (D) HaloTag TMR ligand was used to label newly synthesized (C) or pre-existing (D) Halo-ATR protein. Left, cell lysates were subjected to immunoblotting with the indicated antibodies. TMR signal was measured by direct detection of fluorescence. Band intensity was quantified by ImageJ. Right, quantification of TMR relative to α-actinin and normalized to the signal at the 1 h (C) or 0 h (D) timepoint. Data is presented as mean ± s.d. (n=3 independent experiments). Results of unpaired t-test between −dTAG and +dTAG conditions are shown (**p<0.01; n.s., not significant). Asterisk besides GNB1L immunoblots indicate non-specific bands.

**Figure S10.**
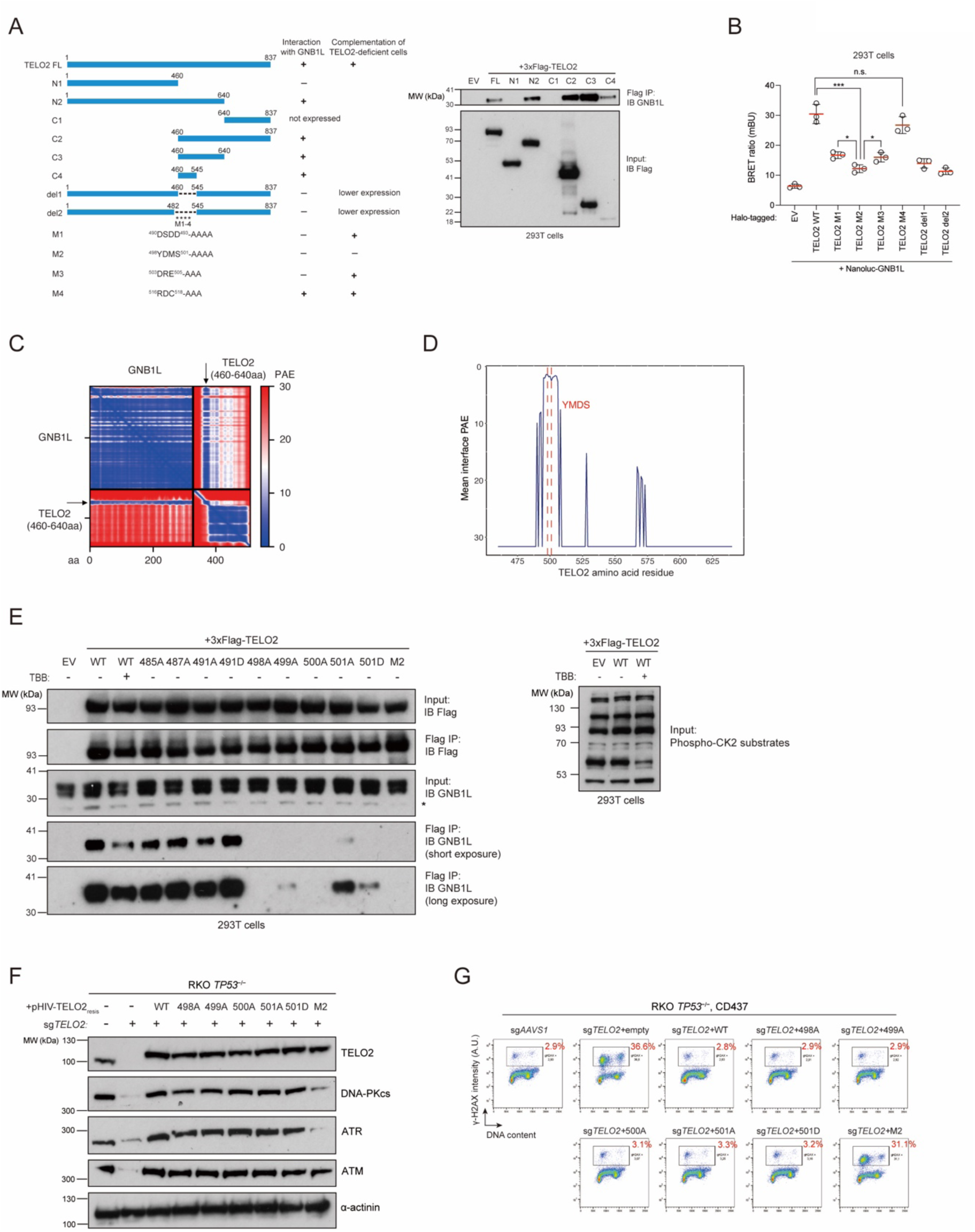
Mapping the GNB1L-TELO2 interaction site on TELO2. Related to Figure 7. (A) Left, schematic of truncation/point mutation mapping in the TELO2 protein and the summary of mutant TELO2 phenotypes. Right, anti-Flag immunoprecipitation (IP) of lysates from 293T cells expressing full-length(FL) or mutant 3xFlag-TELO2. EV, empty vector. The GNB1L-TELO2 interaction was examined by immunoblot analysis with a GNB1L antibody. (B) NanoBRET assay to assess the in vivo interaction between GNB1L and mutant TELO2. Bars represent the mean ± s.d. (n=3 independent experiments). Comparisons were made using an unpaired t-test (***p<0.001; *p<0.05; n.s., not significant). (C) The predicted aligned error (PAE) matrix of GNB1L and the TELO2 aa460-640 fragment, generated by AlphaFold2-multimer. The off-diagonal quadrants represent inter-chain PAE scores. The blue stripes indicated with arrows reflect the set of residues that have a good confidence of relative inter-chain position. (D) Plot of the mean predicted aligned error between each TELO2 residue and GNB1L residues within 9 Å. The four residues ^498^YMDS^501^ are highlighted by red dashed lines. (E) Left, anti-Flag immunoprecipitation (IP) in 293T cells expressing 3xFlag-tagged wildtype (WT) or mutant TELO2. One sample expressing WT TELO2 was treated with 75 μM of the CK2 inhibitor TBB for 20 hours. The GNB1L-TELO2 interaction was examined by immunoblotting with a GNB1L antibody. Right, CK2 inhibition by TBB was confirmed by immunoblot analysis with an antibody recognizing phospho-CK2 substrates. EV, empty vector. (F), (G) RKO *TP53^−/−^* cells were infected with lentiviruses expressing sg*TELO2* and sgRNA-resistant TELO2 variant constructs as indicated. WT, wildtype. (F) Immunoblot analysis of cell lysates with the indicated antibodies. α-actinin, loading control. (G) Cells were treated with 250 nM CD437 for 24 hours, and then fixed, stained with a γ-H2AX antibody and DAPI. Red numbers indicate the percentage of γ-H2AX positive cells. A.U., arbitrary units.

**Figure S11.**
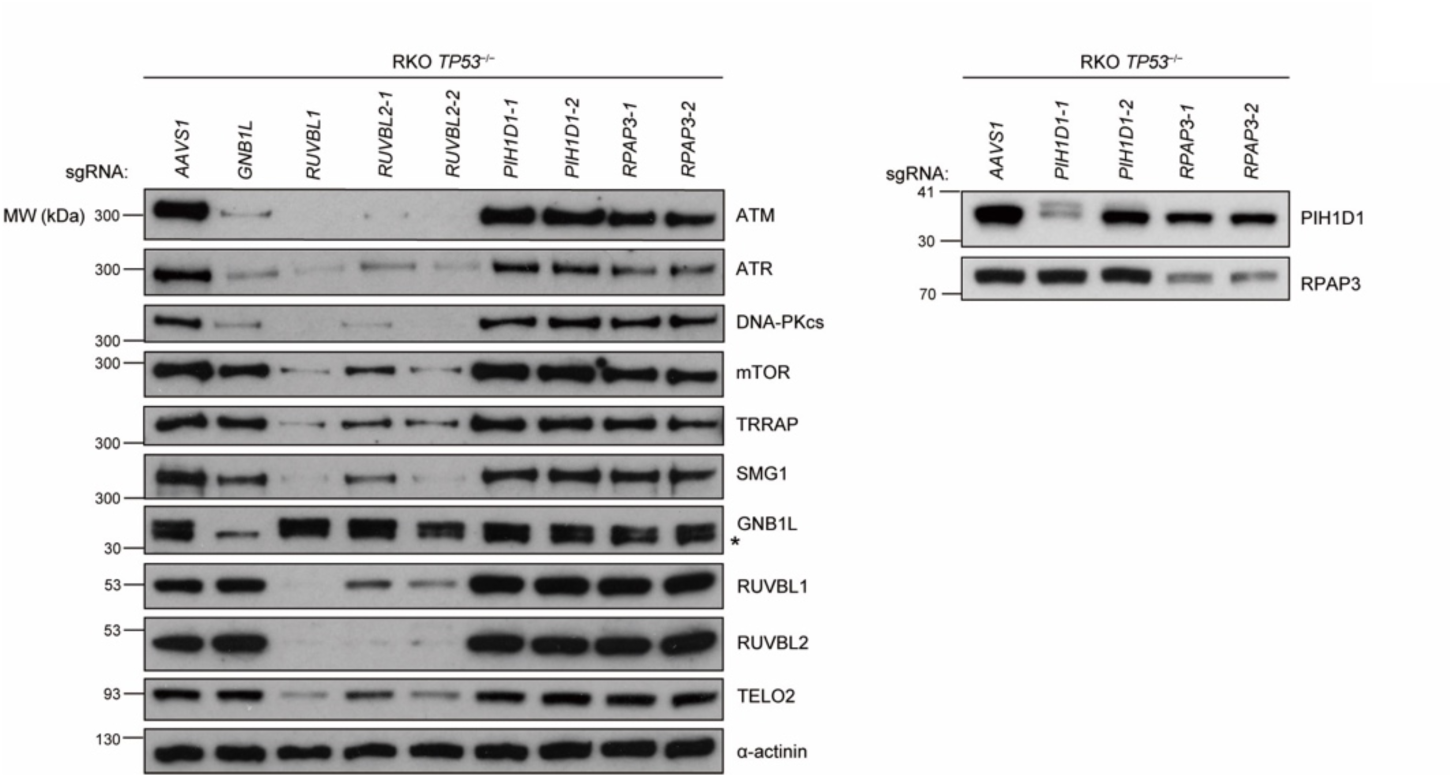
*PIH1D1* and *RPAP3* are not essential to maintain PIKK protein levels in RKO *TP53^−/−^* cells. RKO *TP53^−/−^* cells were transduced with lentiviruses expressing the indicated sgRNA and cell lysates were subjected to immunoblot analysis. α-actinin, loading control. Asterisk besides GNB1L immunoblots indicate non-specific bands.

